# Terpenes from *Cannabis sativa* Induce Antinociception in Mouse Chronic Neuropathic Pain via Activation of Spinal Cord Adenosine A_2A_ Receptors

**DOI:** 10.1101/2023.03.28.534594

**Authors:** Abigail M. Schwarz, Attila Keresztes, Thai Bui, Ryan J. Hecksel, Adrian Peña, Brianna Lent, Zhan-Guo Gao, Martín Gamez-Rivera, Caleb A. Seekins, Kerry Chou, Taylor L. Appel, Kenneth A. Jacobson, Fahad A. Al-Obeidi, John M. Streicher

## Abstract

Terpenes are small hydrocarbon compounds that impart aroma and taste to many plants, including *Cannabis sativa*. A number of studies have shown that terpenes can produce pain relief in various pain states in both humans and animals. However, these studies were methodologically limited and few established mechanisms of action. In our previous work, we showed that the terpenes geraniol, linalool, β-pinene, α- humulene, and β-caryophyllene produced cannabimimetic behavioral effects via multiple receptor targets. We thus expanded this work to explore the efficacy and mechanism of these *Cannabis* terpenes in relieving chronic pain. We first tested for antinociceptive efficacy by injecting terpenes (200 mg/kg, IP) into male and female CD- 1 mice with chemotherapy-induced peripheral neuropathy (CIPN) or lipopolysaccharide-induced inflammatory pain, finding that the terpenes produced roughly equal efficacy to 10 mg/kg morphine or 3.2 mg/kg WIN55,212. We further found that none of the terpenes produced reward as measured by conditioned place preference, while low doses of terpene (100 mg/kg) combined with morphine (3.2 mg/kg) produced enhanced antinociception vs. either alone. We then used the adenosine A_2A_ receptor (A_2A_R) selective antagonist istradefylline (3.2 mg/kg, IP) and spinal cord-specific CRISPR knockdown of the A_2A_R to identify this receptor as the mechanism for terpene antinociception in CIPN. *In vitro* cAMP and binding studies and *in silico* modeling studies further suggested that the terpenes act as A_2A_R agonists. Together these studies identify *Cannabis* terpenes as potential therapeutics for chronic neuropathic pain, and identify a receptor mechanism in the spinal cord for this activity.

## Introduction

Chronic pain is a debilitating condition that causes suffering and decreases patient quality of life, which impacts 20.5% of the United States adult population [1]. Opioids are not effective for treating many chronic pain types, including limited efficacy in neuropathic pain, and opioids are further burdened by adverse side effects, such as constipation, addiction, and tolerance [2–5]. Consequently, patients have increasingly turned to *Cannabis sativa* to treat various conditions, including chronic pain [6]. *Cannabis* has been used in traditional medicinal practices for thousands of years, and recent legalization and access have increased its use in the United States [7]. The primary cannabinoids cannabidiol (CBD) and Δ-9-tetrahydrocannabinol (THC) have been shown to be effective in managing chronic pain in some studies. However, their efficacy is generally moderate, and THC is burdened by unwanted psychoactive side effects [8, 9]. These limits have focused attention on other potentially therapeutic components of *Cannabis*, including minor cannabinoids, flavonoids, and terpenes.

*Cannabis* is unique in the number of terpenes it contains; while most other plants have two dominating terpene species, *Cannabis* contains up to 150 terpenes, with multiple terpenes acting as the dominant species [10]. This complexity of the *Cannabis* chemovar may determine the different biological effects caused by different strains of *Cannabis* [11]. A number of studies in both humans and animals have suggested that terpenes can relieve pain, including chronic neuropathic pain [12]. However, these studies are limited by methodological constraints, like a lack of critical controls, the use of complex undefined terpene mixtures, limited translational testing (e.g. side effects), and small sample sizes, along with a near-total lack of investigation into the mechanism by which terpenes produce pain relief [12].

To fill this gap, we investigated the terpenes α-humulene, β-caryophyllene, β-pinene, geraniol, and linalool, which are found in moderate to high levels within *Cannabis*. We found that these terpenes have cannabimimetic properties, namely hypothermia, hypolocomotion, catalepsy, and antinociception [13]. Notably, the antinociceptive effects of these terpenes for acute pain were blocked by the Cannabinoid Receptor 1 (CBR1) antagonist rimonabant, suggesting a CBR1 mechanism for antinociception. Furthermore, we showed that each terpene activated CBR1 *in vitro* [13]. Some behaviors were not inhibited by blocking CBR1, but were blocked by treatment with the Adenosine A_2A_ Receptor (A_2A_R) antagonist istradefylline, suggesting multiple receptor involvement in the mechanism of these *Cannabis* terpenes [13].

Moving forward from this work, we sought to investigate whether these terpenes are effective in a chronic pain model and to determine their mechanism of action using *in vivo*, *in vitro*, and *in silico* methods. Our findings suggest that all five terpenes in this study act on the spinal cord A_2A_R to elicit pain relief in chronic neuropathic pain, potentially at the receptor’s orthosteric site. By shedding light on the therapeutic potential of *Cannabis* terpenes, this study contributes to the search for more effective treatments for chronic pain with fewer side effects, and fills the gap in the literature around the effects and mechanisms of terpenes in chronic pain relief.

## Methods

### Drugs

The terpenes α-humulene (Tokyo Chemical Industry, Lot #MTQZC-AJ), β-caryophyllene (Tokyo Chemical Industry, Lot #ARY4A-HD), β-pinene (Alfa Aesar, Lot #10217611), geraniol (Alfa Aesar, Lot #10211653), and linalool (Alfa Aesar, Lot #10216518) were diluted in 10% DMSO, 10% Tween80 and 80% USP Saline for injections. The small molecule inhibitors rimonabant (ApexBio, Lot #B1429-2) and istradefylline (Tokyo Chemical Industry, Lot #2PJUO-MT) were dissolved in ratios of 10:10:80 and 20:10:70 in DMSO:Tween80:USP saline, respectively. Morphine sulfate pentahydrate was obtained from the National Institute on Drug Abuse Drug Supply Program and was dissolved in USP saline. WIN55,212-2 (Sigma-Aldrich, Lot #0000026263) was diluted in 10% DMSO, 10% Tween80 and 80% USP saline for injections. Lipopolysaccharide (LPS; Enzo, Lot #07282012) was diluted in USP saline. Paclitaxel (Thermo Scientific, Lot #Q22I058) was used to induce Chemotherapy-Induced Peripheral Neuropathy (CIPN). The drug was dissolved in 15:15:70 cremophor-EL (Fisher Scientific, #23-847-01SET):ethanol:USP Saline and administered via intraperitoneal (IP) injection at 2mg/kg. Doxorubicin (Fisher Scientific, #BP251610), NECA (Fisher Scientific, #16-911-0), and CP55,940 (Sigma Aldrich, #C1112-10MG) were only used for *in vitro* experiments, prepared as below. All solutions were made fresh for each experiment and used immediately. Matched vehicle injections were included in each experiment as a control. For *in vitro* experiments, 100 mM stock solutions of the terpenes in DMSO were made, and 10 mM stock solutions in DMSO of all other compounds. All final solutions were made at 1% DMSO with matched vehicle controls.

### Animals

Male and female CD-1 (a.k.a. ICR) mice aged 5-8 weeks from Charles River Laboratories were used for all experiments. The mice were acclimated to the temperature and humidity-controlled vivarium at the University of Arizona for at least five days after arrival. 12-hour light and dark cycles were maintained, and standard chow and water were available *ad libitum*. Animals were acclimated to the experiment room for 30 minutes before any procedure. The mice were randomly assigned to treatment groups, and the experimenters were blinded to treatment groups through coded drug vials; unblinding only occurred once all data for each experiment was collected. The IACUC at the University of Arizona approved all experiments. Experiments were also carried out in accord with the NIH Guide for the Care and Use of Laboratory Animals, and the International Association for Pain Guidelines for the Use of Animals in Research.

### Chemotherapy-Induced Peripheral Neuropathy (CIPN)

This model was induced and measured for mechanical allodynia as in our previous work [14]. Paclitaxel (taxol) was used to induce CIPN. Animals were baselined for mechanical sensitivity using von Frey filaments using the up-down method as in [15] and in our previous work [14] and then injected with 2 mg/kg IP paclitaxel on days 1, 3, 5, and 7 of the experiment. On day 8, the animals had their post-treatment baseline measured to ensure they had developed neuropathy, and then were given various drugs over various treatment regimens and routes of administration (IP, SC, IT, PO) as described in the Figure Legends. Mechanical sensitivity was measured for a three-hour time course.

### Acute Inflammatory Model

Male and female mice were baselined for mechanical sensitivity using von Frey filaments as above. The animals received an intraplantar (IPL) injection into their left hind paw of 100 ng of LPS to induce inflammatory pain. Their mechanical threshold was measured again 10 minutes later to insure that inflammatory pain was established. They were then given either an injection of terpene (200 mg/kg, IP) or vehicle 20 min after LPS injection, and mechanical sensitivity was measured for three hours, after which the mice were sacrificed. Their hind paw skin tissue was collected for cytokine analysis.

### Conditioned Place Preference (CPP)

CPP was performed as in our previous work [16]. Male and female CD-1 mice were labeled and weighed each day. Mice were brought to the testing room to acclimate for at least 30 minutes before further handling by the researcher. On Day 0, mice were baselined and allowed to freely explore all chambers of the 3-chambered CPP apparatus (Spatial Place Preference LE 896/898 rigs), and their placement was recorded for 15 minutes. On Days 1-4, mice were injected twice daily with either drug treatment (terpene, 200mg/kg IP) or vehicle (10% DMSO, 10% Tween-80, 80% USP saline) and placed into their respective paired or unpaired chamber according to a counter-balanced schedule (vehicle one time, terpene the other, varying between morning and afternoon). Groups were injected at either 9 am and 1 pm or 10:15 am and 2:15 pm. On day 5, conditioned mice could freely explore the CPP apparatus and were placed into the center hallway chamber. Their placement was recorded for 15 minutes, then compared to the baseline. Data was recorded and analyzed by PPC WIN 2.0 software.

### Cannabinoid Tetrad Testing

Tetrad testing was performed similarly to our previous work [13]. The mice had baseline measurements taken for tail flick (52°C water, 10 sec cutoff), body temperature (rectal thermometer), and catalepsy (ring stand test, immobile/cataleptic behavior over 5 min). Mobile time baselines in the open field test were not measured, since we previously showed baseline testing produces significant locomotor habituation [13]. Mice then received either drug treatment (terpene, 200-500 mg/kg PO or vaporization as below) or vehicle (10% DMSO, 10% Tween80, 80% USP saline). Ten minutes following treatment, the open field test was performed by placing mice in a square plastic arena [13] and recording with a video camera for 5 minutes. Locomotor video data was analyzed using ANY-Maze software. This was followed directly by the catalepsy ring test, with immobile/cataleptic behavior recorded for 5 min. Tail flick thermal nociception was then assessed at 20 min post-treatment. Finally, body temperature for each mouse was recorded at 30 min post-treatment. For some experiments, the tail flick time course was extended past this point to 120 min. Under this paradigm, each single mouse was assessed for all four tetrad behaviors.

### Terpene Inhalation by Vaporizer

Baseline tetrad measurements were first recorded prior to terpene exposure as described above. Mice were then placed in individual compartments attached to a Storz and Bickel Volcano Hybrid Vaporizer with vacuum tubing. Terpenes were vaporized at or near their respective boiling points: 198°C for linalool, 230°C for geraniol, 167°C for β-pinene, and 230°C for β-caryophyllene. The vaporizer was loaded with 2 mL of pure terpene and animals were exposed for 5 minutes. This exposure interval was repeated for a total of 4 exposures (8 mL terpene for 20 minutes). Catalepsy, tail flick, and body temperature were recorded at 5, 10, and 20 min post-exposure, respectively, as described above.

### CRISPR Knockdown

This approach was also performed as in our previous work [17]. All-in-one predesigned CRISPR constructs with both gRNA and Cas9 expression cassettes were purchased from Genecopoeia. The negative control (NC) CRISPR construct contained identical elements to the knock-down CRISPR construct but with a non-targeting gRNA (#CCPCTR01-CG01-B). The knock-down construct targeted the *Adora2a* (Adenosine A_2A_ Receptor) mouse gene (#MCP290877-CG12-3-B). All CRISPR DNA constructs were given at a dose of 750 ng DNA in a 5 µL volume with GeneJet *in vivo* transfection reagent (#SL100500, Lot# 8051662) by the manufacturer’s protocol and delivered via intrathecal (IT) injection as described in our previous work [14, 18]. Mice were given 2 injections daily for 3 days, at least 3 hours apart. Behavior experiments and tissue collection occurred on day 10. Day 3 of CRISPR injections coincided with day 1 of paclitaxel injections in the CIPN model above, so that day 10 of CRISPR and day 8 of CIPN coincided.

### Immunohistochemistry (IHC)

IHC was performed essentially as reported in our previous work [18]. Negative control and A_2A_R knockdown CRISPR mice were perfused with saline followed by 4% paraformaldehyde, and their spinal cords were collected immediately after the behavioral experiments’ completion. The spinal cords were fixed in 4% paraformaldehyde (PFA) and soaked in 15% and then 30% sucrose in phosphate buffered saline solution (PBS), pH 7.4. Tissue samples were embedded and frozen in OCT medium at -20°C. 30 µm sections of the spinal cord were sliced via cryostat and immediately underwent preparation for IHC. Tissue sections were soaked for 30 minutes at room temperature (RT) in endogenous peroxidase block (60% methanol, 0.3% H_2_O_2_, 39.7% PBS), followed by blocking buffer (5% normal donkey serum, 1% bovine serum albumin in PBS with 0.1% Tween20 [PBST]) for one hour at RT and left overnight in the A_2A_R primary antibody at 4°C (Abcam #ab3461, Lot # GR3431019-2, 1:100) in blocking solution. The secondary antibody goat anti-rabbit Alexa488 (Invitrogen #A11034, Lot# 2286890, 1:500) in the same solution was incubated at RT for 1 hr, washed with PBST, then preserved in Fluormount and imaged using an Olympus BX51 microscope equipped with a Hamamatsu C8484 digital camera. Images were analyzed using ImageJ. Whole spinal images were selected, and average mean intensities were measured, background was subtracted, then compared between groups.

### Cell Culture

For A_2A_R molecular pharmacology experiments, we used a HEK293 cell line stably expressing the human A_2A_R [19]. The cells were cultured in Dulbecco’s Modified Eagle Medium (DMEM, Gibco) supplemented with 10% heat-inactivated fetal bovine serum (Gibco), 1% penicillin-streptomycin, and 2 µmol/mL of glutamine. For *in vitro* inflammatory mechanism experiments, we used a mouse microglial BV-2 cell line expressing an NFκB-luciferase reporter; this was a kind gift from Dr. Valeri Mossine from the University of Missouri [20]. The BV-2 cells were cultured in DMEM supplemented with 5% heat-inactivated fetal bovine serum (Gibco) and 1% penicillin-streptomycin. Both lines were cultured in a humidified incubator at 37°C with 5% CO_2_ and sub-cultured every 2-3 days.

### In Vitro Assay – cAMP Accumulation

A_2A_R-HEK cells were grown for 24 hrs in 96 well plates. The medium was then removed, and the cells washed twice with PBS. Cells were then treated with assay buffer containing the A_2B_R antagonist PSB603 (1 µM, Tocris), rolipram (30 µM, Sigma) and adenosine deaminase (3 U/ml, Worthington) for 30 min followed by addition of agonists or vehicle control and incubated for 20 min. The reaction was terminated by removal of the mixture of assay buffer and compounds, and addition of 50 µl 0.3% Tween-20. cAMP levels were measured with an Amplified Luminescent Proximity Homogeneous Assay (ALPHA) Screen cAMP assay kit as instructed by the manufacturer (PerkinElmer, Waltham, MA).

### In Vitro Assay – Competition Radioligand Binding

A_2A_R binding inhibition procedures were similar to our previously reported methods [19]. Briefly, membrane preparations from A_2A_R-HEK cells were incubated at 25°C for 60 min with a range of terpene concentrations in the presence of the selective radioligand [3H]ZM241385 (1.0 nM, American Radiolabeled Chemicals). The incubation was terminated by rapid filtration with vacuum using an MT-24 cell harvester (Brandell, Gaithersburg, MD, USA) with GF/B filters (Whatman), and the filters were washed twice rapidly with chilled Tris−HCl buffer (∼5 mL). Scintillation counts were measured using a Tri- Carb2810TR counter (PerkinElmer). The resulting data was normalized to the percent competition caused by vehicle (0%) vs. saturating non-radioactive ligand (100%).

### In Vitro Assay - NF-kB Luciferase Reporter

15,000 cells/well of the BV-2 NFκB reporter cells were plated in a 96 well plate 24 hours prior to experimental use. Cells were then serum starved (DMEM media alone) for 1 hour, pre-treated with 500 µM of individual terpene, 10 µM WIN55,212, or matched vehicle for 5 minutes, after which 1 µg/mL of LPS was added for 3 hrs at 37°C. After stimulation, cells were lysed via a 15 minute - 20°C freeze with lysis buffer (10% glycerol, 2 mM diaminocyclohexanetetraacetic acid, 25 mM Tris phosphate, and 1% Triton-X100). The plate was then centrifuged at 4000 RPM at 4°C for 5 minutes. Cell lysate was transferred to a white 96 well plate where it was combined with luciferase substrate (20 mM Tris, 10 mM magnesium chloride, 0.1 mg/mL bovine serum albumin, 0.5 mM ethylenediaminetetraacetic acid, 33.3 mM dithiothreitol, 1 mM ATP, 30 µM sodium pyrophosphate, and 1.1 mg/mL D-luciferin). Luminescence was obtained at 560-580 nm with a 10-minute read delay via CLARIOstar Plus plate reader.

### In Vitro Assay - Resazurin Cell Viability Assay

15,000 cells/well of the BV-2 cells were plated in a black 96 well plate 24 hours prior to experimental use. Cells were gently washed with PBS to remove phenol red media. 500 µM terpene, 10 µM doxorubicin, 10 µM CP55,940, 10 µM WIN55,212, or matched vehicle (0.5% DMSO) were added to appropriate wells and incubated for 1 hour at 37°C. 0.075 mg/mL resazurin (Fisher Scientific, #AC41890-0050) in PBS was added to each well and incubated for a further 2 hours at 37°C. Fluorescence was quantified via a CLARIOstar Plus plate reader using 550 nm excitation and 590 nm emission.

### In Silico *Modeling*

All terpene docking interactions with the A_2A_R and the mu opioid receptor (MOR) were studied by molecular docking simulations using the general docking method available in Molecular Operating Environment (MOE) 2022.02. Ligand geometry was determined by energy minimization and charge correction as recommended by MOE. After loading the PDB files of the A_2A_R (PDB:3QAK) and MOR (PDB:4DAK) crystal structures, the protein structures were subjected to the Structure Preparation application as outlined in MOE. The structure preparation is used to prepare 3D biomolecular structures for docking simulations. The molecular features of the terpenes (**Table S1**) were opened in MOE and the DOCK activated. The receptor protein structure opened in MOE and each terpene structure was subjected to 100 poses using London dG followed by 5 poses for refining under GBVI/WSA dG. After completion of the docking, the data was analyzed by selecting the refined 5 poses of each ligand and combined with the receptor in MOE to generate the structures of the receptor-ligand complex. To view the interactions between the ligand and receptor the protein contact was activated and analyzed based on binding energy and distance of interacting groups between each ligand pose and the receptor. Pairwise energy of interactions between the receptor amino acids and the ligands of < -1.0 Kcal/Mol and a distance of < 4.0 Å were applied to select the most interacting residues.

### Cytokine Analysis by qPCR

qPCR was performed as reported in our previous work [21]. Flash-frozen paw skin was homogenized, and RNA was extracted using Trizol and chloroform from the ipsilateral (LPS treated) and contralateral (no treatment control) hind paws of CD-1 mice post-behavior analysis. The Trizol manufacturer’s protocol was followed. 1 µg of total RNA was converted to cDNA via reverse transcription using a Reverse Transcriptase cDNA Kit (Fisher Scientific, #43-874-06) by the manufacturer’s protocol. The cytokines IL-6 (F: 5’-TCC AGT TGC CTT CTT GGG AC-3’ R: 5’-GTG TAA TTA AGC CTC CGA CTT G-3’), IL-10 (F: 5’- GAC CAG CTG GAC AAC ATA CTG CTA A-3’ R: 5’-GAT AAG GCT TGG CAA CCC AAG TAA-3’), and TNF-α (F: 5’-CTC TTC AAG GGA CAA GGC TG-3’ R: 5’-TGG AAG ACT CCT CCC AGG TA-3’) were amplified from the cDNA samples using PowerUp SYBR Green Master Mix (Thermo-Scientific, Lot #01284919). The housekeeping gene GAPDH (F: 5’TCC TGC ACC ACC AAC TGC TTA G-3’ R: 5’-GAT GAC CTT GCC CAC AGC CTT G-3’) was used to control for loading. Cycle thresholds were measured, normalized to GAPDH, and transformed by 1/2^cycledifference, then directly compared to measure mRNA quantity.

### Data Analysis

All data were reported as the mean ± SEM, and in a few cases, normalized to a control group as described in the appropriate Figure Legends. The behavioral data were reported raw, without Maximum Possible Effect or other normalization (with the exception of the WIN55,212 dose/response curve analysis). Biological and technical replicates are described in the Figure Legends. Area Under the Curve (AUC) was calculated for many of the behavioral time course data sets; this was done using the AUC function in GraphPad Prism 9.5 with the pre-pain baseline data point excluded from analysis. Behavioral time course comparisons were performed using a Repeated Measures (RM) 2 Way ANOVA with a Sidak’s (2 groups; e.g. LPS pain) or a Dunnett’s (3 groups or more; e.g. terpene + morphine combination) *post hoc* test. AUC comparisons were performed depending on their experimental design. Analysis from single experiments (e.g. morphine + terpene combination in CIPN) were performed by 1 Way ANOVA with Tukey’s *post hoc* test. Combined analysis from separate experiments (e.g. LPS pain) was performed by 1 Way ANOVA with Fisher’s Least Significant Difference *post hoc* test; this uncorrected test is justified by the independent nature of each comparison, established by separate experiments. For AUC analysis of antagonist treatment (e.g. istradefylline in CIPN), each pair was compared by an unpaired 1-Tailed *t* Test, justified since each experiment was performed separately as a comparison set, with a significant effect of antagonist on the time course data, justifying a 1-Tailed hypothesis. CPP analysis was performed by a RM 2 Way ANOVA with a Sidak’s *post hoc* test, with each conditioned group compared to their own pre-conditioning baseline. The *in vitro* experimental results were analyzed by a 1 Way ANOVA with Dunnett’s *post hoc* test. Comparison of 2 single groups (e.g. vaporizer experiment) was performed by an unpaired 2-Tailed *t* Test. In all cases, significance was defined as p < 0.05. Linear regression to calculate the A_50_ of WIN55,212 in CIPN was performed as described in our previous work [22]. All graphing and statistical analysis was performed using GraphPad Prism 9.5. Males and females were included in every experiment and compared by 2 Way ANOVA. No sex differences were detected, so all males and females were combined for analysis.

## Results

### Terpenes are Efficacious in Relieving Neuropathic Pain

As noted above, we found that the terpenes geraniol, linalool, α-humulene, β-pinene, and β- caryophyllene had modest efficacy in relieving acute nociceptive tail flick pain [13]. We thus expanded from this work to test their efficacy in relieving mechanical allodynia in a model of chemotherapy-induced peripheral neuropathy (CIPN). As a comparison control and to establish dosing, we first tested the synthetic non-selective cannabinoid WIN55,212 for efficacy in this model (1-10 mg/kg, IP). We found that WIN55,212 produced efficacious, time- and dose-dependent antinociception that peaked at or above pre-CIPN baseline levels (**Figure 1A**). Dose/response analysis revealed an A_50_ for WIN55,212 of 1.5 mg/kg (**Figure 1B**).

**Figure 1:**
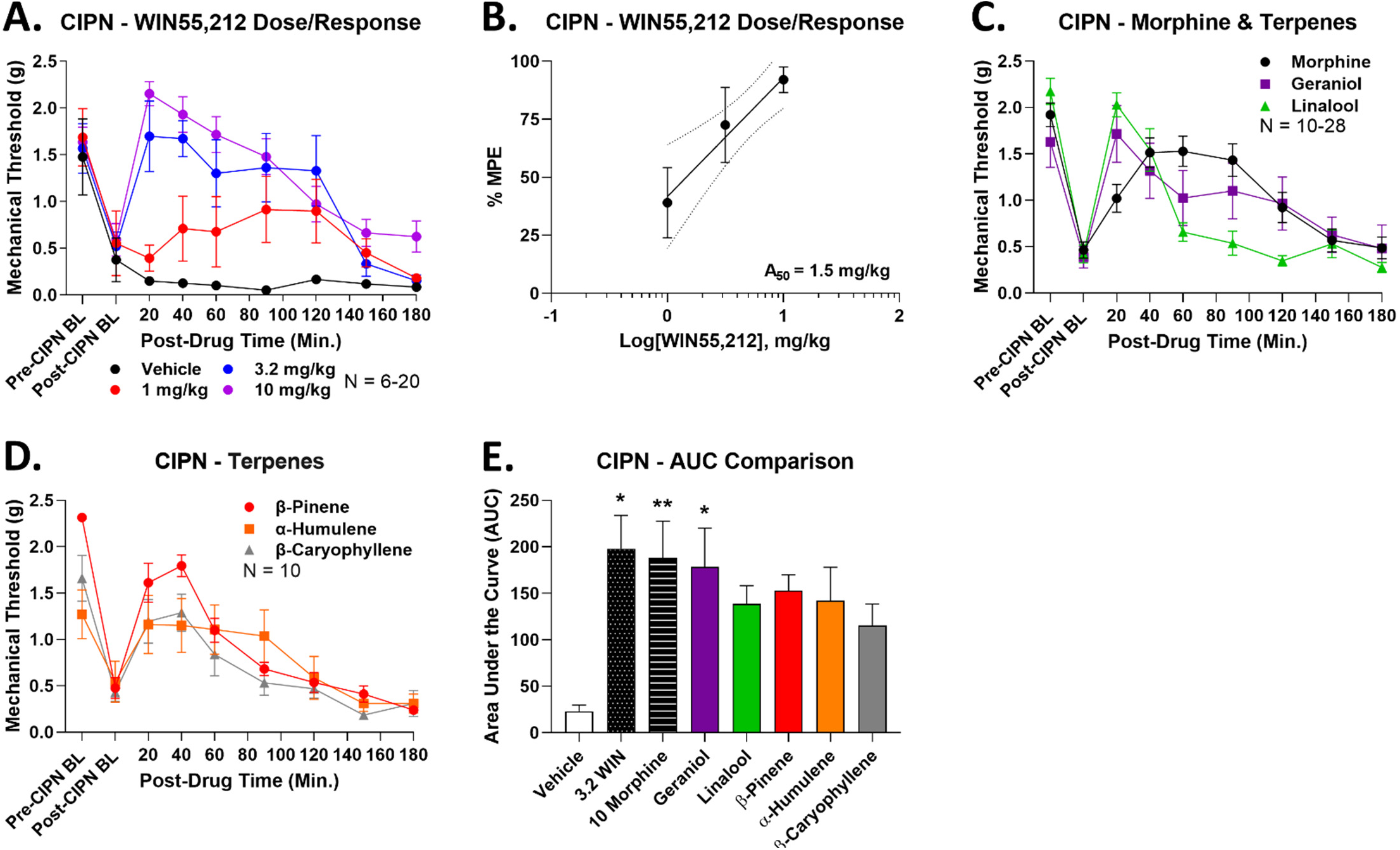
Terpenes are efficacious in relieving neuropathic pain. Male and female CD-1 mice had CIPN induced and measured as described in the Methods. Data shown is the mean ± SEM, performed in 2-6 technical replicates for each experiment, with sample sizes noted in each graph. BL = baseline. **A)** WIN55,212 or vehicle injected (1-10 mg/kg, IP). Each dose had a vehicle comparison performed at the same time, which are all combined here. **B)** The data from **A** was normalized to %Maximum Possible Effect (%MPE) and used to construct a dose/response curve. Linear regression revealed an A_50_ = 1.5 mg/kg for WIN55,212 in CIPN. The dotted lines are the 95% confidence intervals for the regression. **C-D)** Each terpene (200 mg/kg, IP) or morphine comparison (10 mg/kg, SC) was injected. Each experiment had a morphine comparison, which are all combined here. **E)** The AUC for each terpene along with vehicle, morphine, and 3.2 mg/kg WIN55,212 controls are shown here. *, ** = p < 0.05, 0.01 vs. vehicle control by 1 Way ANOVA with Fisher’s Least Significant Difference *post hoc* test.

Following this, we tested each terpene individually (200 mg/kg, IP, dosing from [13]) along with a morphine comparison (10 mg/kg, SC) for efficacy. Each terpene also produced efficacious time-dependent antinociception near or above the peak effect of morphine (**Figure 1C-D**). Area Under the Curve (AUC) analysis of this time course data showed that 3.2 mg/kg WIN55,212, 10 mg/kg morphine, and 200 mg/kg geraniol all produced significant antinociception over vehicle control (**Figure 1E**). Although the other terpenes did not have a statistically significant elevation, their AUC mean was still 5-6 fold elevated over the vehicle mean and close to the mean values for WIN55,212, morphine, and geraniol (**Figure 1E**). Taken together, this evidence suggests that all terpenes produce robust pain relief in CIPN.

### Terpenes are Efficacious in Relieving Inflammatory Pain

To expand our analysis of terpene antinociceptive efficacy, we next tested for terpene pain relief in a model of lipopolysaccharide (LPS)-induced inflammatory pain. Mechanical allodynia was produced by LPS in all mice (**Figure 2A**). Most terpenes (200 mg/kg, IP) produced significant time-dependent antinociception over vehicle control; the only exception was β-pinene, which produced a small, non-significant improvement in mechanical threshold (**Figure 2A**). AUC analysis backed up this conclusion, with geraniol and linalool both producing significant elevation in AUC over vehicle control (**Figure 2B**). Much like with CIPN above, while the other terpenes had non-significant AUC increases, the mean values were still elevated 5-7 fold over the vehicle mean (**Figure 2B**). Both data types together suggest that all terpenes except β-pinene are effective antinociceptive agents in this second, different pathological pain type.

**Figure 2:**
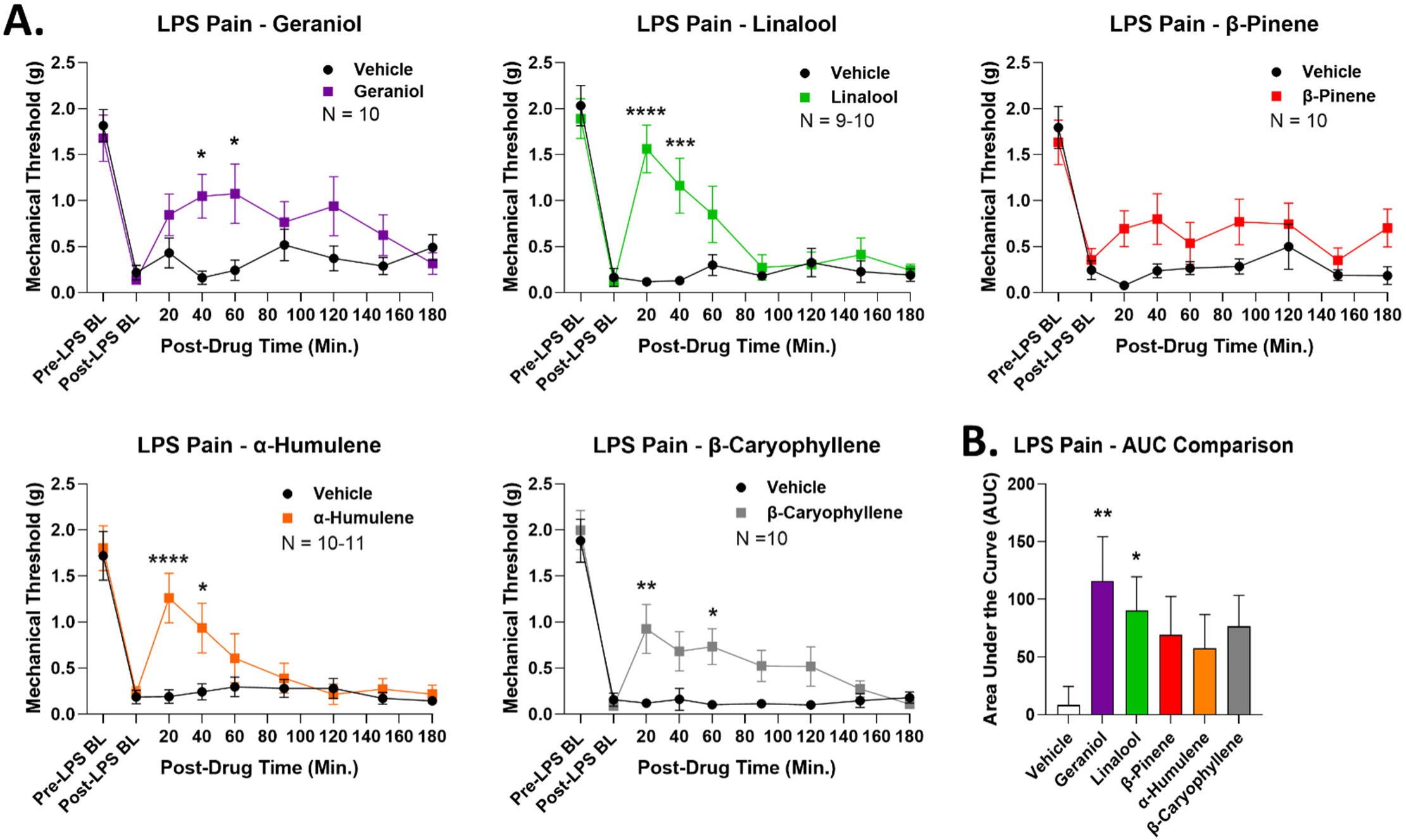
Terpenes are efficacious in relieving acute inflammatory pain. Male and female CD-1 mice had inflammatory pain induced by LPS and measured as described in the Methods. Data shown is the mean ± SEM, performed in 2 technical replicates for each experiment, with sample sizes noted in each graph. BL = baseline. **A)** Terpene (200 mg/kg, IP) injected as noted along with vehicle control. *, **, ***. **** = p < 0.05, 0.01, 0.001, 0.0001 vs. same time point vehicle group by RM 2 Way ANOVA with Sidak’s *post hoc* test. **B)** AUC values calculated for each terpene and vehicle control, shown here. All vehicle results were combined into the group shown. *, ** = p < 0.05, 0.01 vs. vehicle group by 1 Way ANOVA with Fisher’s Least Significant Difference *post hoc* test.

### Terpenes Have No Reward Liability

Terpenes have been shown to produce pain relief in various models in both humans and animals [12]. However, there has been very little exploration of other crucial translational features of terpene therapy, including side effects. We thus sought to investigate the potential reward liability of our terpenes using conditioned place preference (CPP), which has only been tested for a few terpenes in a limited way, and mostly for their impact on other drugs of abuse [23–26]. We tested for terpene reward liability in an unbiased, counter-balanced 4-day CPP conditioning protocol (see Methods).

As expected, vehicle treatment showed neither preference nor aversion, while morphine (10 mg/kg) showed a positive preference, validating the assay (**Figure 3A**). Crucially, both geraniol and linalool showed neutral conditioning, neither preference nor aversion, suggesting they do not cause reward or dysphoria (**Figure 3A**). When combined with our pain data above, this suggests these terpenes could be effective analgesics with no rewarding or dysphoric side effects. In contrast, both α-humulene and β-caryophyllene showed a significant place aversion, suggesting they may be dysphoric under these treatment conditions (**Figure 3A**). β-pinene did not show a significant difference vs. pre-conditioning baseline; however, the post-conditioning mean was still in the aversive direction, suggesting potential aversive/dysphoric side effects. Before-and-after analysis of each single mouse backed up this group analysis (**Figure 3B**). Overall, these results suggest that no terpene has reward liability, some have neither reward nor aversive liability, while some have aversive liability. These observations are crucial when evaluating the potential clinical utility of these ligands as pain drugs.

**Figure 3:**
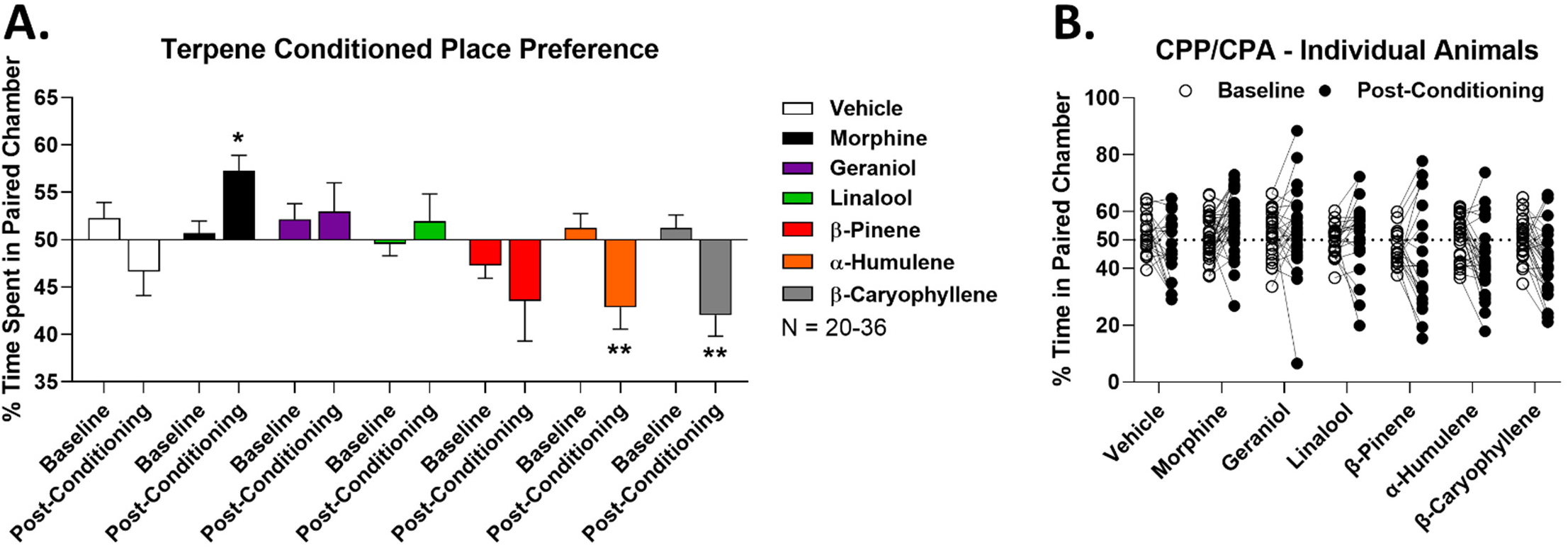
Terpenes lack reward liability as measured by conditioned place preference. Male and female CD-1 mice had vehicle, morphine (10 mg/kg, SC), or terpene (200 mg/kg, IP) injected over a 4 day conditioning trial (see Methods), with preference measurement on day 5. Data shown represents the mean ± SEM of the % time spent in paired chamber over 3-5 technical replicates per group, with sample sizes noted in the legend. Values over 50% represent preference while values under 50% represent aversion. **A)** Summary data shown for each group. *, ** = p < 0.05, 0.01 vs. the baseline for each group by RM 2 Way ANOVA with Sidak’s *post hoc* test. Morphine shows a positive preference (reward), validating the assay. **B)** The baseline and post-conditioning trajectories for each mouse in each group are shown.

### Terpenes Enhance Morphine Antinociception in CIPN

Analgesic additivity/synergy is an important translational feature of some drugs, whereby lower doses of 2 analgesics enhance each other’s pain relief efficacy while not enhancing side effects, leading to an overall improved therapeutic profile. As an example, some CB2-selective cannabinoids where shown to synergize with morphine for pain relief, while leading to lower side effects for both drug classes [27]. In addition, in our earlier work we showed that all terpenes in this study produced enhanced pain relief in tail flick pain when combined with the synthetic cannabinoid WIN55,212 [13]. We thus tested for the ability of our terpenes to produce enhanced pain relief in CIPN when combined with other analgesics.

We first combined a lower dose of terpene (100 mg/kg, IP) with a lower dose of morphine (3.2 mg/kg, SC) in CIPN as above. We show full time course curves for geraniol in **Figure 4A**, and all other full time course data is in **Figure S1**. All terpenes showed the same pattern. The separate lower doses of morphine and terpene produced measurable but modest antinociception in CIPN, which in all cases were statistically the same as each other. When combined, *all* terpene/morphine combinations showed a significant elevation over either terpene or morphine alone at a minimum of 3 separate time points (**Figures 4A, S1**). AUC analysis solidified this conclusion, with the AUC of all terpene/morphine combination treatments significantly elevated over single drug treatment (**Figure 4B**). This data suggests that terpenes could be combined with opioids in CIPN pain as a means to improve the therapeutic index of both treatments.

**Figure 4:**
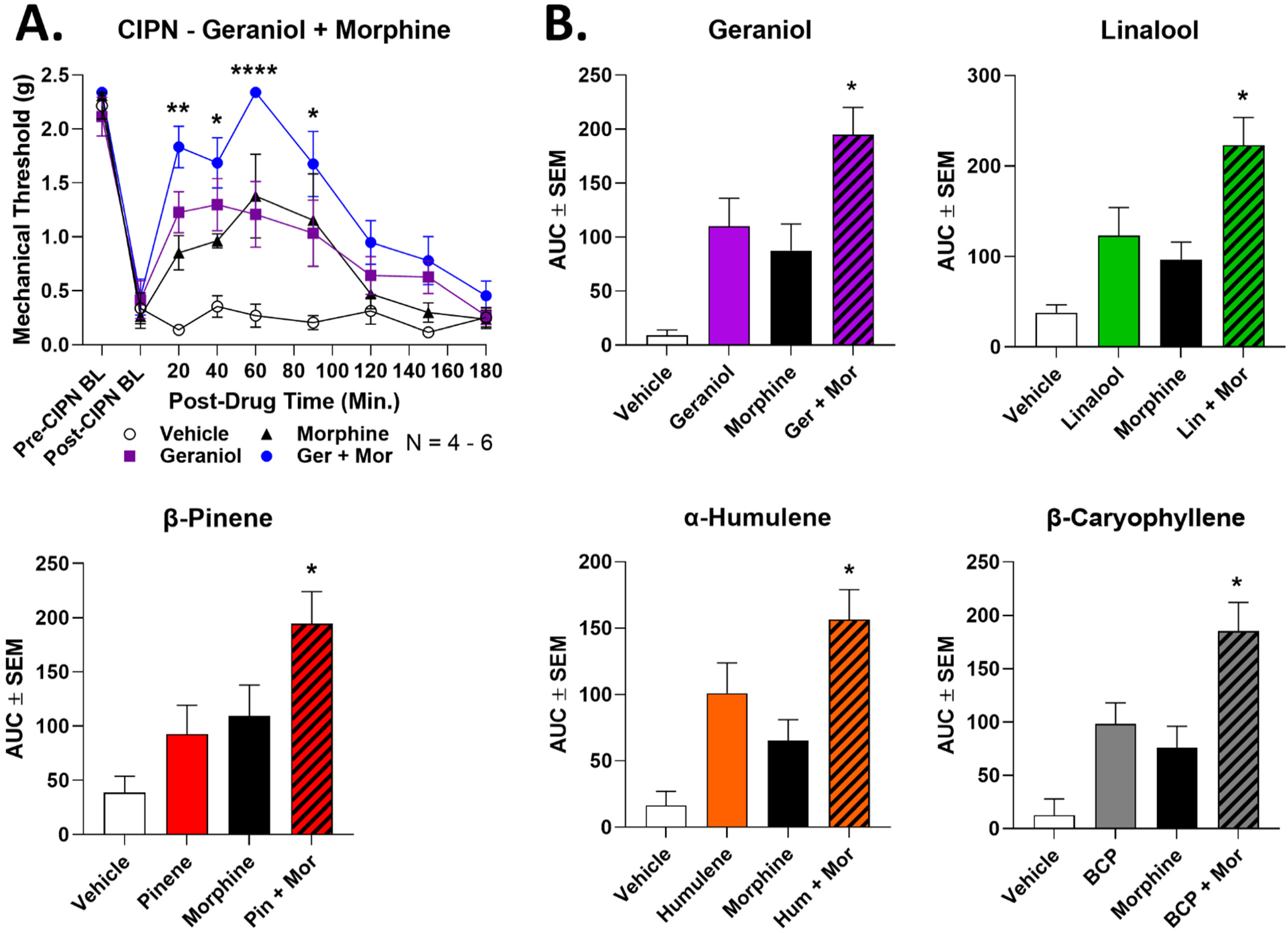
Terpenes enhance morphine pain relief in CIPN. Male and female CD-1 mice had CIPN induced and measured as described in the Methods. Data shown is the mean ± SEM, performed in 2-3 technical replicates for each experiment, with sample sizes noted in each graph. BL = baseline. Vehicle, morphine (3.2 mg/kg, SC), terpene (100 mg/kg, IP) or both combined were injected, with mechanical allodynia measured. **A)** The time course data for geraniol is shown as an example; the curves for the other terpenes are all shown in **Figure S1**. *, **, **** = p < 0.05, 0.01, 0.0001 vs. same time point morphine or terpene group by RM 2 Way ANOVA with Dunnett’s *post hoc* test. **B)** The AUC data from **A** was calculated and is shown here. * = p < 0.05 vs. terpene or morphine group by 1 Way ANOVA with Tukey’s *post hoc* test.

Based on our earlier work, we also tested a lower dose of terpene (100 mg/kg, IP) with a lower dose of WIN55,212 (1 mg/kg, IP) in CIPN. Unlike with morphine, terpene and WIN55,212 combination produced either a weak or non-existent enhancement over either alone. Geraniol, β-pinene, and β-caryophyllene did have a significant elevation of terpene/WIN55,212 treatment at a single time point during their time course (**Figure S2**). However, AUC analysis did not show a significant elevation of any terpene/WIN55,212 combination over either single treatment (**Figure S3**). Taken together, this suggests that our terpenes do not effectively enhance WIN55,212 antinociception in CIPN, unlike morphine above, and unlike our earlier work in tail flick pain [13]. Overall, these results suggest that terpene/morphine but not terpene/cannabinoid combinations could be effective in managing CIPN.

### Terpenes Produce Comparable Antinociceptive Tolerance to Morphine in CIPN

Another key translational feature of pain therapies is antinociceptive tolerance with repeated dosing, whereby a drug becomes less effective at the same dose over time as the system adapts and activates negative feedback loops. Notably, we could not find any studies testing this for terpenes. We thus established a twice-daily 4-day treatment regimen in CIPN with daily mechanical threshold measurements, testing morphine (10 mg/kg, SC) and each terpene (25-200 mg/kg, IP). The AUC analysis of each dose over the 4-day regimen is shown in **Figure 5**. All raw time course data for each drug and dose is shown in **Figures S4-S6**.

**Figure 5:**
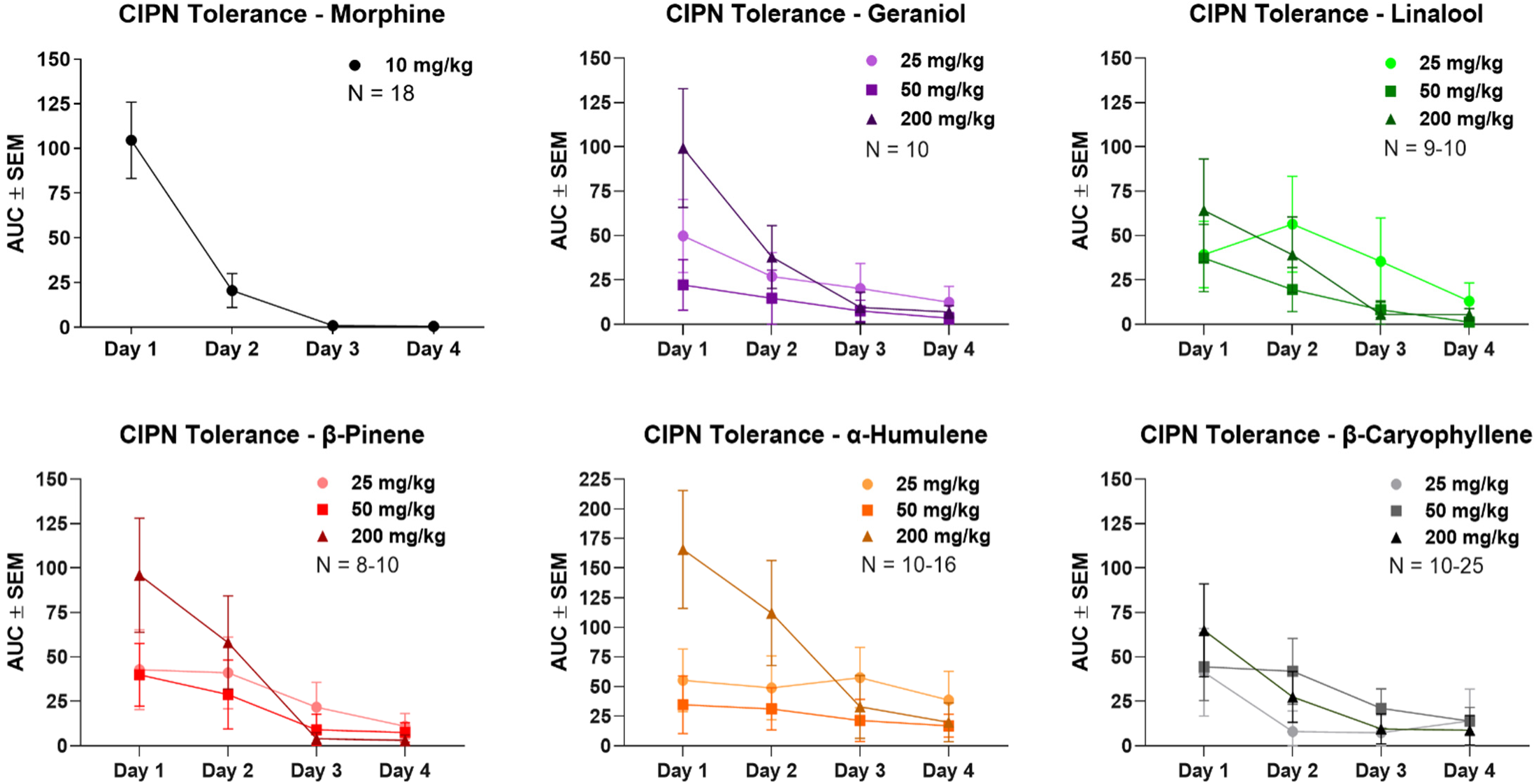
Terpenes produce similar antinociceptive tolerance to morphine in CIPN. Male and female CD- 1 mice had CIPN induced as in the Methods, with morphine (10 mg/kg, SC) or terpene (25-200 mg/kg, IP) injection twice daily over a 4-day protocol, with daily mechanical allodynia measurement after the morning injection. Data shown is the mean ± SEM, performed in 2-5 technical replicates for each experiment, with sample sizes noted in each graph. The data shown here is the AUC calculated from each experimental set, with all raw data shown in **Figures S4-S6**. The morphine data is reproduced with permission from [21]. The 200 mg/kg dose of each terpene showed a roughly similar tolerance trajectory to 10 mg/kg morphine, with perhaps a less severe drop off on day 2. Lower doses of terpene had less initial efficacy, but also appear to have had slower tolerance development.

Under this treatment paradigm, morphine showed rapid tolerance in CIPN, with a ∼75% loss in AUC response by day 2 and no detectable pain relief by day 4. Each terpene at 200 mg/kg showed comparable but slightly slower tolerance to morphine, with less than ∼50% loss in response by day 2, but still a total loss of response by day 4. Lower doses of terpene (25-50 mg/kg, IP) showed less initial efficacy than at 200 mg/kg, but did appear to have slower tolerance development. At lower doses most terpenes showed little day-to-day loss, and all had measurable pain relief at day 3, while 25 mg/kg α-humulene still had a measurable response at day 4. Overall, these data make clear that terpenes do produce antinociceptive tolerance in CIPN with repeated dosing, but this tolerance is no worse than and may be better than the established therapeutic morphine.

### Terpenes Have Limited Bioavailability by the Oral and Inhalation Routes of Administration

Our studies above used the IP injection route to deliver terpenes. However, this route is not very translationally relevant, and the pharmacokinetics of terpenes have not been tested in the literature to our knowledge. We thus performed preliminary route of administration studies to test terpene bioavailability by relevant routes. We first tested a limited set of terpenes by the oral route of administration (gavage, PO) at 200 mg/kg. We used the cannabinoid tetrad as an output measure since most of the behaviors are highly sensitive and we know the terpenes produce activity in these assays [13]. This dose and route produced no measurable tail flick antinociception or locomotor changes, but did cause small but significant hypothermic effects for geraniol and β-pinene (**Figure S7A**). From these observations we quickly concluded that terpenes had limited bioavailability at this dose and route, so we increased the dose to 500 mg/kg, and tested the full set of terpenes and controls. At this higher dose, we still could not detect changes in tail flick antinociception (**Figure S7B**). However, geraniol did produce a small but significant decrease in locomotor activity, and all terpenes produced modest but significant hypothermia (**Figure S7B**). This data together suggests that terpenes have limited bioavailability by this route.

In addition to oral dosing, we tested the inhalation route by exposing mice to vaporized pure terpene for 20 minutes in a vaporization chamber (see Methods). All terpenes tested produced modest but significant hypothermia (**Figure S8A**). However, they did not effectively increase tail flick antinociception; only β- caryophyllene had a significant elevation, but the effect size was very small (**Figure S8B**). Lastly, none of the vaporized terpenes caused an increase in catalepsy (**Figure S8C**). This data also suggests that terpenes have limited bioavailability by this route. Future studies will need to explore means to improve terpene pharmacokinetics in order to maximize translatability.

### Terpenes Produce Antinociception in CIPN via the Adenosine A_2A_ Receptor in Spinal Cord

Apart from therapeutic evaluation, we also sought to identify a mechanism of action for our terpenes in CIPN, as very few such mechanistic studies have been performed [12]. In our earlier work, we found that the A_2A_R antagonist istradefylline was able to block some of the tetrad behaviors evoked by terpene treatment [13]. We thus used istradefylline to test for A_2A_R engagement by terpenes in CIPN. First, we performed control experiments to establish a non-confounding dose of istradefylline, as this drug had motor activating effects in our hands [13]. At 10 mg/kg IP, istradefylline caused a significant decrease in mechanical thresholds in naïve mice, ruling out this dose (**Figure S9A**). However, 3.2 mg/kg IP istradefylline had no impact on mechanical thresholds in naïve mice, suggesting this dose was valid for experimental use (**Figure 6A**). We also tested this dose of istradefylline alone in CIPN, which had no impact on mechanical thresholds in mice undergoing this pain state (**Figure S9B**).

**Figure 6:**
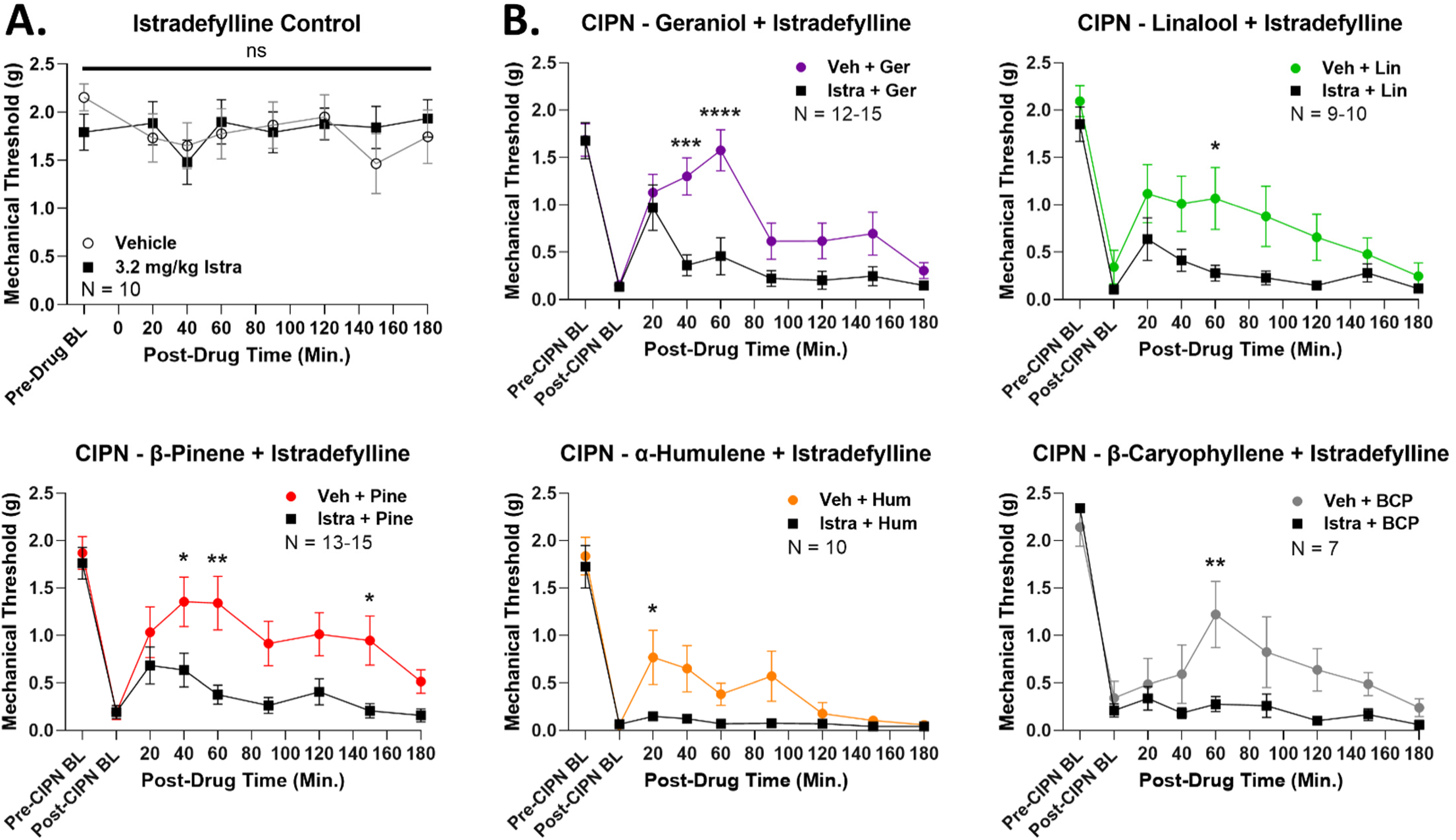
Terpenes evoke antinociception in CIPN via the A_2A_R. Male and female CD-1 mice had CIPN induced and measured as described in the Methods. Data shown is the mean ± SEM, performed in 2-3 technical replicates for each experiment, with sample sizes noted in each graph. BL = baseline. **A)** Naïve mice with no CIPN had vehicle or the A_2A_R antagonist istradefylline (3.2 mg/kg, IP) injected, followed by mechanical threshold measurements. Istradefylline had no impact on naïve mechanical thresholds, validating the dose. **B)** CIPN mice had vehicle or istradefylline (3.2 mg/kg, IP) injected, with a 10 min treatment time, followed by terpene (200 mg/kg, IP) as noted. *, **, ***, **** = p < 0.05, 0.01, 0.001, 0.0001 vs. same time point istradefylline/terpene group by RM 2 Way ANOVA with Sidak’s *post hoc* test. Istradefylline significantly reduced antinociception in CIPN by each terpene, implicating the A_2A_R as a mechanism of action.

Thus validated, we used 3.2 mg/kg istradefylline to test for A_2A_R engagement by terpenes in CIPN. Istradefylline treatment caused a significant decrease in antinociception for all terpenes in CIPN for at least one time point in their time course (**Figure 6B**). This was further validated by AUC analysis, which showed a significant decrease caused by istradefylline pre-treatment for each terpene (**Figure 7**). This data strongly suggests that terpenes produce antinociception in CIPN at least in part by activating the A_2A_R.

**Figure 7:**
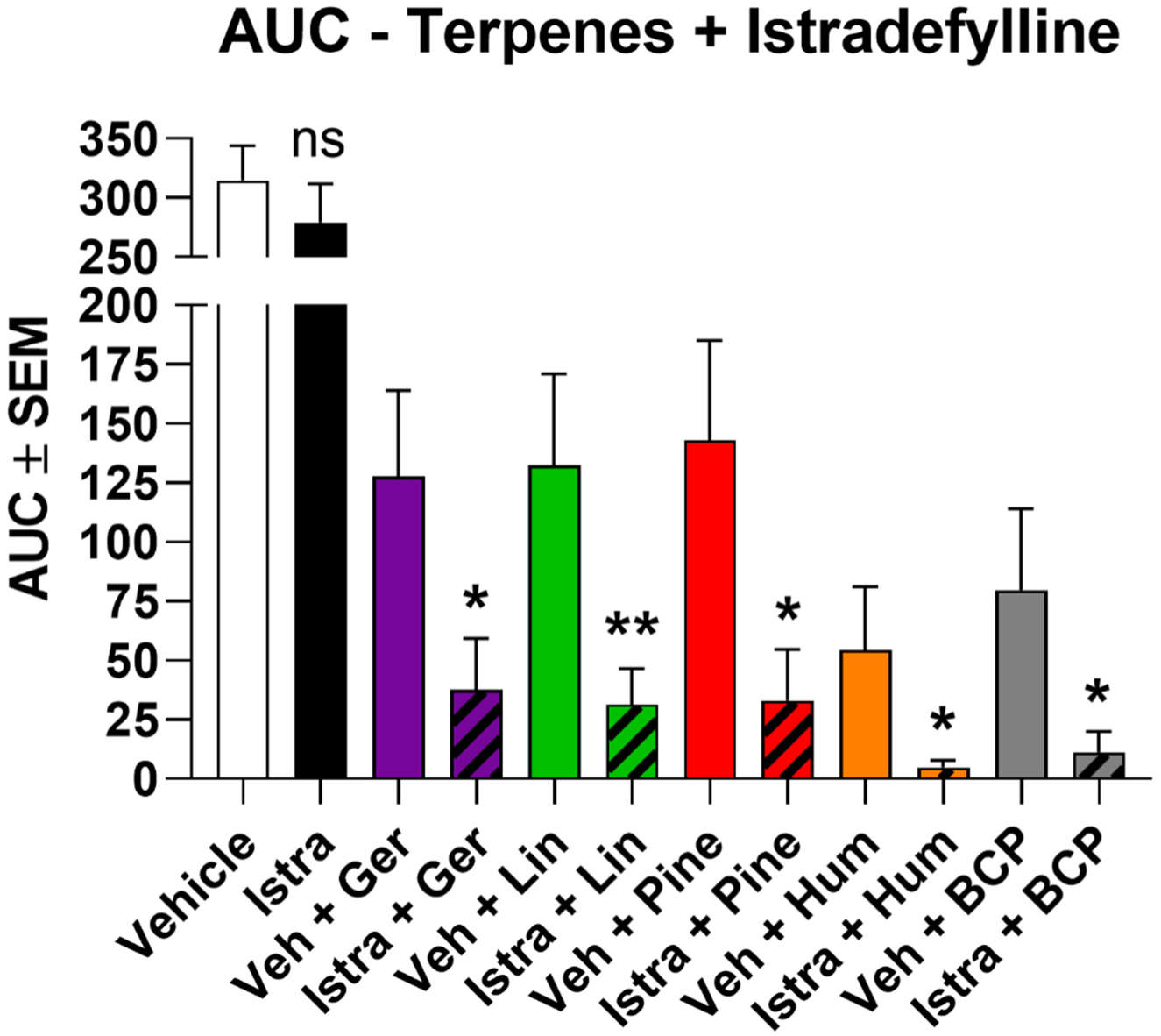
AUC data for istradefylline/terpene treatment in CIPN. The AUC was calculated from each pair of experiments in Figure 6 and shown here. *, ** = p < 0.05, 0.01 vs. paired vehicle/terpene group by Unpaired 1- Tailed *t* Test. The comparison is justified by the independent nature of each experiment along with the demonstrated reduction in antinociception with istradefylline treatment shown in Figure 6.

Since our earlier work showed that terpenes evoke tail flick anti-nociception via the CBR1 [13], we also used the CBR1 antagonist rimonabant to test for CBR1 engagement in CIPN. We first validated a 10 mg/kg, IP dose of rimonabant, showing it had no effect on mechanical thresholds alone (**Figure S10A**). We then tested each terpene, finding in contrast to our earlier work that rimonabant had no impact on terpene time courses (**Figure S10B**) or AUC values (**Figure S10C**) in CIPN. Our data thus suggested that the A_2A_R but not the CBR1 is a mechanism for terpene pain relief in CIPN.

Up to this point, all of our drugs were given by systemic routes, meaning that sites of action could be anywhere in the body. To narrow down a terpene site of action, we performed intrathecal injections of terpene directly into the spinal cord during CIPN (100 nmol, IT). By this route, all terpenes except for β-pinene caused significant antinociception over vehicle control, confirmed by both time course (**Figure 8A**) and AUC (**Figure 8B**) analysis. This experiment suggested that most terpenes could act via the spinal cord to produce pain relief in CIPN, although this does not rule out other additional sites of action.

**Figure 8:**
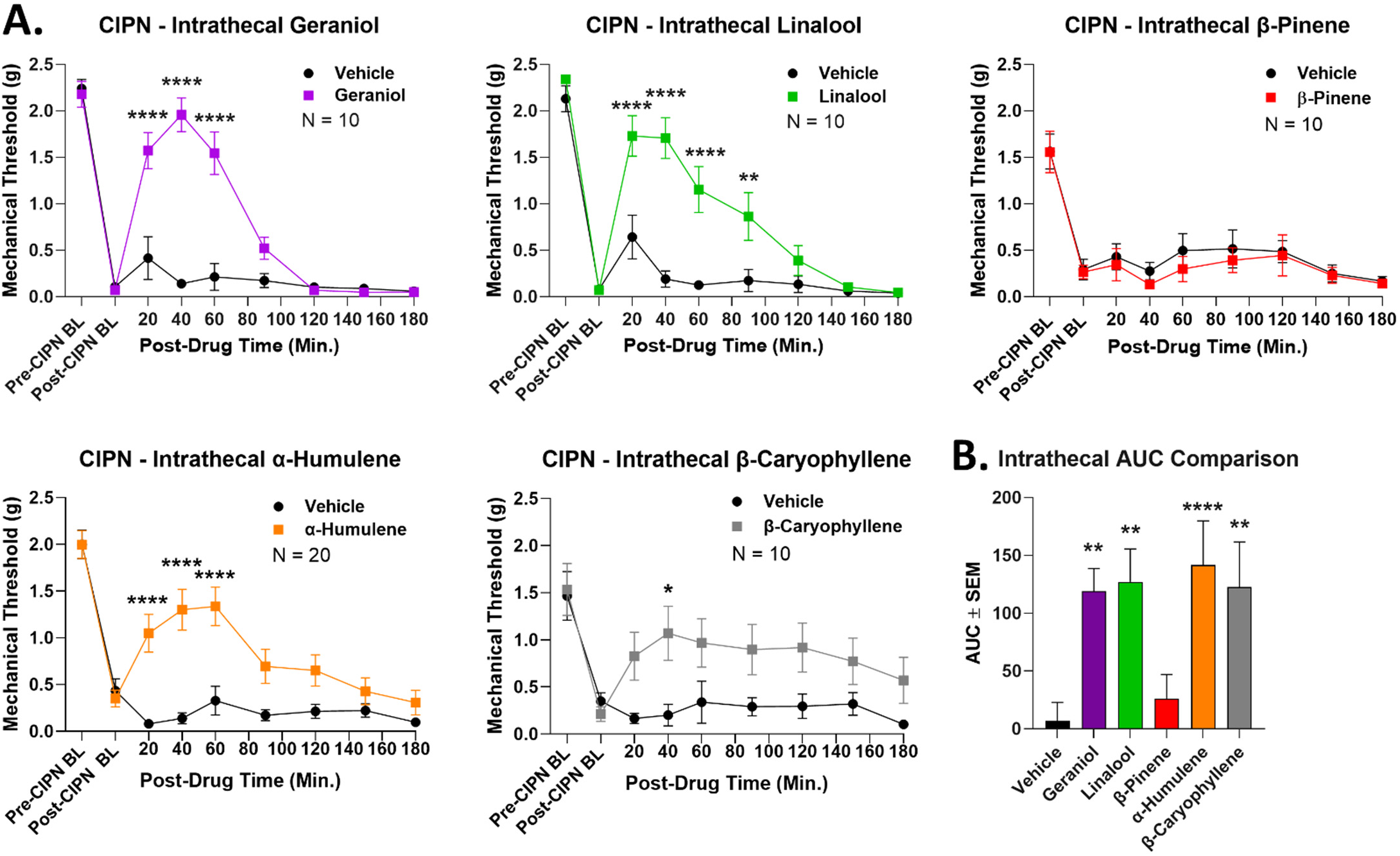
Most terpenes evoke antinociception in CIPN via a spinal cord site of action. Male and female CD-1 mice had CIPN induced and measured as described in the Methods. Data shown is the mean ± SEM, performed in 2-4 technical replicates for each experiment, with sample sizes noted in each graph. BL = baseline. **A)** Vehicle or terpene (100 nmol, IT) injected into the spinal cord in CIPN mice and mechanical allodynia measured. *, **, **** = p < 0.05, 0.01, 0.0001 vs. same time point vehicle group by RM 2 Way ANOVA with Sidak’s *post hoc* test. **B)** AUC values calculated from the data in **A** (all vehicle groups combined). **, **** = p < 0.01, 0.0001 vs. vehicle group by 1 Way ANOVA with Fisher’s Least Significant Difference *post hoc* test. All terpenes except β-pinene induce antinociception when injected into the spinal cord.

Once we had identified an A_2A_R mechanism and a spinal cord site of action, we connected these two findings by performing CRISPR-mediated knockdown of the A_2A_R in the spinal cord. This was achieved by repeated IT injection of CRISPR DNA construct into adult wild type mice, allowing for acute receptor knockdown during CIPN. This treatment caused a significant decrease in terpene antinociception in at least one time point in each time course (**Figure 9A**). β-pinene was not tested since the results above suggest a non-spinal cord site of action. AUC analysis confirmed a significant decrease with A_2A_R CRISPR treatment for α-humulene and β- caryophyllene, with a near-significant (p = 0.07) decrease for linalool (**Figure 9B**). Geraniol AUC was not significantly decreased; comparing the time course and AUC data, our results suggest that geraniol could be partially but not fully mediated by spinal cord A_2A_R (**Figure 9B**). We also validated successful A_2A_R knockdown in spinal cord using immunohistochemistry (**Figure S11**). Overall, these experiments suggest a mechanism by which most terpenes activate the A_2A_R in spinal cord to produce pain relief in CIPN.

**Figure 9:**
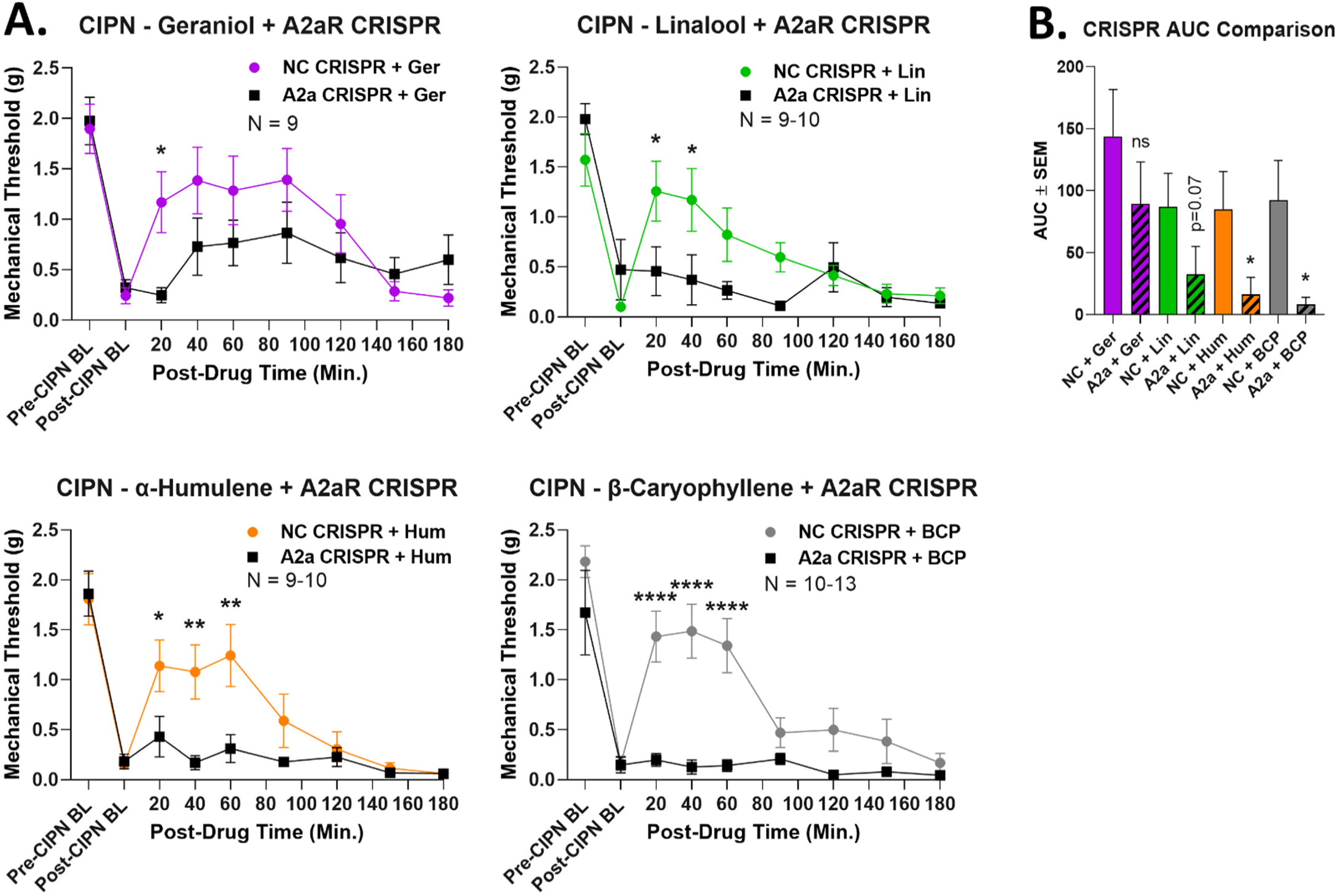
Terpene mechanism confirmed by A_2A_R CRISPR knockdown in the spinal cord. Male and female CD-1 mice had CIPN induced along with A_2A_R-targeted CRISPR or Negative Control (NC) CRISPR injections into the spinal cord so that day 8 of CIPN and day 10 of CRISPR coincided (see Methods). Data shown is the mean ± SEM, performed in 2-3 technical replicates for each experiment, with sample sizes noted in each graph. BL = baseline. Terpene injected (200 mg/kg, IP), with the exception of β-pinene due to lack of intrathecal response above, followed by mechanical allodynia measurement. **A)** Time course data shown. *, **, **** = p < 0.05, 0.01, 0.0001 vs. same time point A_2A_R-CRISPR group by RM 2 Way ANOVA with Sidak’s *post hoc* test. **B)** AUC data calculated from **A** and compared. * = p < 0.05 vs. same terpene NC group by Unpaired 1-Tailed *t* Test. The comparison is justified by the independent nature of each experiment along with the demonstrated reduction in antinociception with A_2A_R-CRISPR treatment shown in **A**.

### Terpenes are Adenosine A_2A_ Receptor Agonists

While the above experiments suggest that these terpenes activate the A_2A_R to produce pain relief, they do not demonstrate whether that is by direct or indirect activation. We thus tested for the ability of the terpenes to directly activate the human A_2A_R in an *in vitro* model system. By using a cAMP accumulation assay, we showed that a 100 µM concentration of each terpene was able to activate the A_2A_R to ∼40-50% of the stimulation produced by the positive control agonist NECA (**Figure 10A**). Notably, this concentration was in the same range or less than that required to produce selective CBR1 activation in our previous work [13]. This data suggests that all terpenes are capable of direct agonism of the A_2A_R.

**Figure 10:**
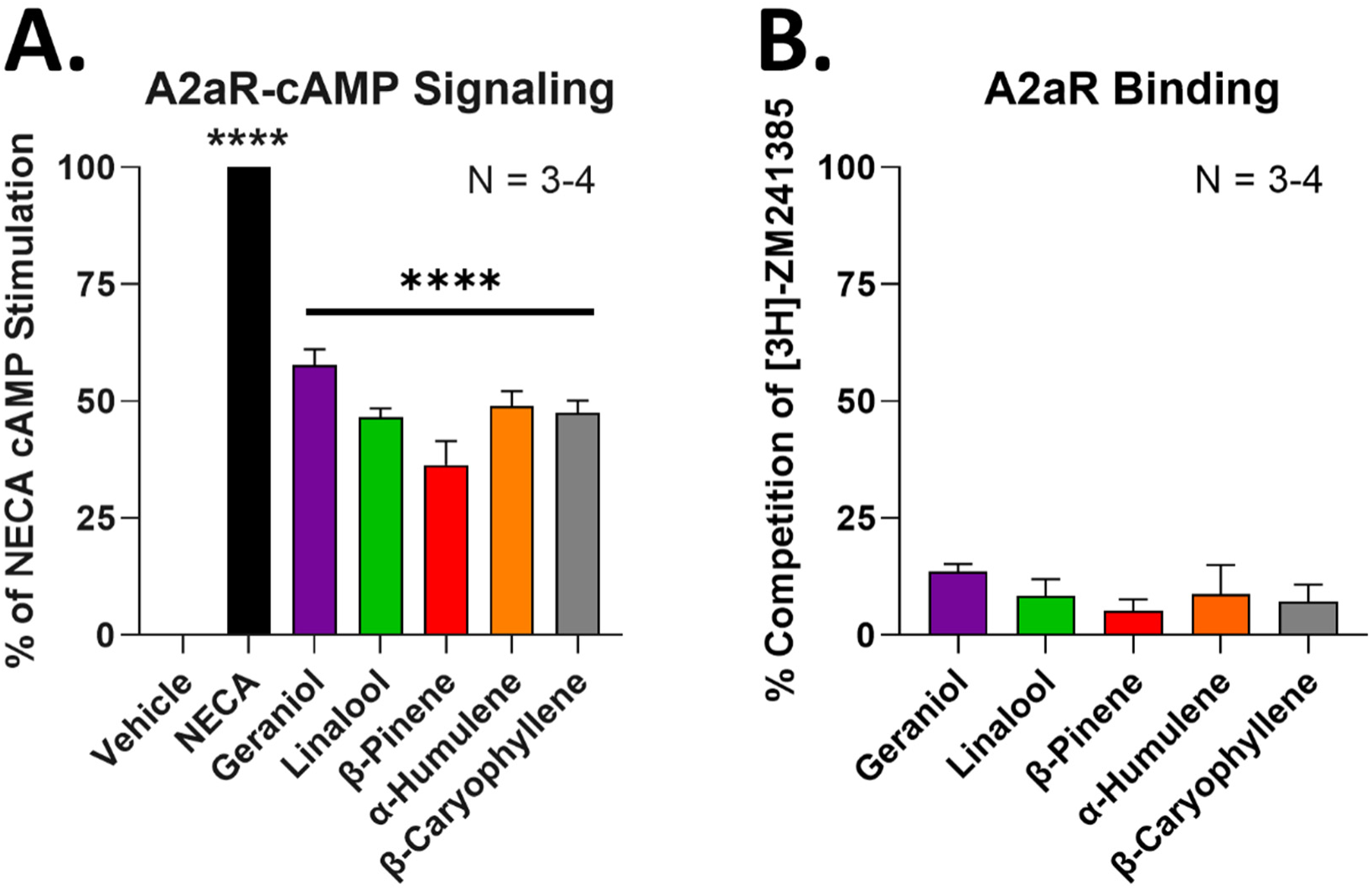
Terpenes act as A_2A_R agonists *in vitro*. A_2A_R-HEK cells used for each experiment. Data represented as the mean ± SEM, with the sample size of independent experiments shown in each graph. Experiments performed in 3-4 technical, independent replicates. **A)** Vehicle, terpenes (100 µM), and NECA positive control (10 µM) used to stimulate cAMP accumulation. Data normalized to the percent stimulation caused by vehicle (0%) and NECA (100%). **** = p < 0.0001 vs. vehicle group by 1 Way ANOVA with Dunnett’s *post hoc* test. **B)** Competition radioligand binding performed with terpene (300 µM) competing against the orthosteric A_2A_R ligand ^3^H-ZM241385. Data normalized to the percent competition caused by vehicle (0%) and saturating cold ligand (100%). Together the results suggest that terpenes act as A_2A_R partial agonists without competing against the orthosteric ligand ZM241385.

In order to investigate further mechanistic details, we performed competition radioligand binding versus the established orthosteric ligand [3H]-ZM241385. Interestingly, none of the terpenes produced effective competition of the radioligand even at a higher 300 µM concentration (**Figure 10B**). This suggests that while the terpenes can activate the receptor, they may do so through a more complex mechanism than straightforward orthosteric agonism, such as allosteric agonism within or outside of the orthosteric binding pocket. This is also similar to our work with these terpenes at the CBR1, where only geraniol produced concentration-dependent radioligand competition at this receptor [13].

In order to provide further insight and potential hypotheses of terpene activation of the A_2A_R, we performed molecular docking simulations of all terpenes at the A_2A_R and at the negative control MOR (no MOR activation by terpene in our previous work [13]). Our initial results suggested very similar docking energy values (Kcal/Mol) for both the A_2A_R and the MOR, which did not explain how the terpenes could activate the A_2A_R without activating the MOR (**Table 1**). However, deeper analysis of the binding poses and active site residue interactions provided potential insight into this question.

**Table 1:**
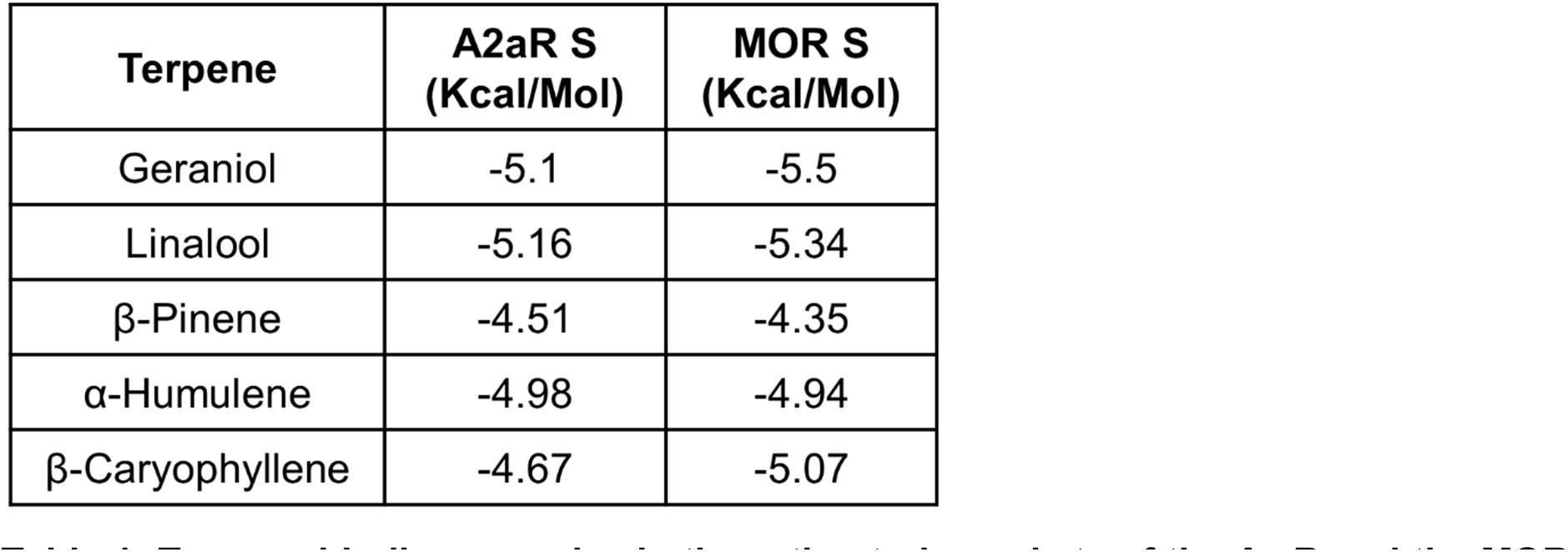
Terpene binding energies in the orthosteric pockets of the A_2A_R and the MOR. Binding energies (Kcal/Mol) as a result of molecular docking simulation are shown for each terpene at each receptor. The binding energies were moderate to low, which may explain the high doses/concentrations required for activation. The energies were also similar for the A_2A_R and the MOR, despite demonstrated lack of terpene activation of the MOR. Insight from the analysis of interacting residues in **Figure 11** may explain this apparent discrepancy.

For the A_2A_R, the docking poses in the orthosteric site and 16 high confidence residue interactions were identified and are shown in **Figure 11A**. There were 8 lipophilic and 8 hydrophilic amino acids. This pattern of 1:1 distribution between nonpolar and polar amino acid side chains is related to the nature of the natural ligand for the A_2A_R, which is a polar compound. Observation of interactions between terpenes and the A_2A_R is driven mainly by hydrophobicity of the terpene ligands which is more than 3 times their polarity (**Table S1**; ASA_H vs. ASA_P). Only geraniol and linalool have hydroxyl polar groups in their structures. The presence of these OH groups enhanced the binding energies for these two ligands to the targeted receptors (**Table 1**). Other interactions were observed between the OH groups of geraniol and linalool. The geraniol OH forms a hydrogen bond with Asn253(TMH-6) and His250(TMH-6). The linalool OH forms a hydrogen bond with Tyr271(TMH-7) and His278(TMH-7). EC2 between TMH-4 and TMH-5 forms part of the binding pocket of the A_2A_R [28] and both Glu169(EC-2) and Phe168(EC-2) form multiple hydrogen bonds with geraniol and linalool side chain hydroxyls. H_2_O 1204 forms a hydrogen bond with the linalool OH group. Phe168(EC-2) is a major recognition amino acid interacting via pi-pi interaction of the phenyl group with the alkene parts of the terpene structures in addition to the lipophilic side chains. These key residue interactions suggest a mechanism by which the terpenes could activate the A_2A_R.

**Figure 11:**
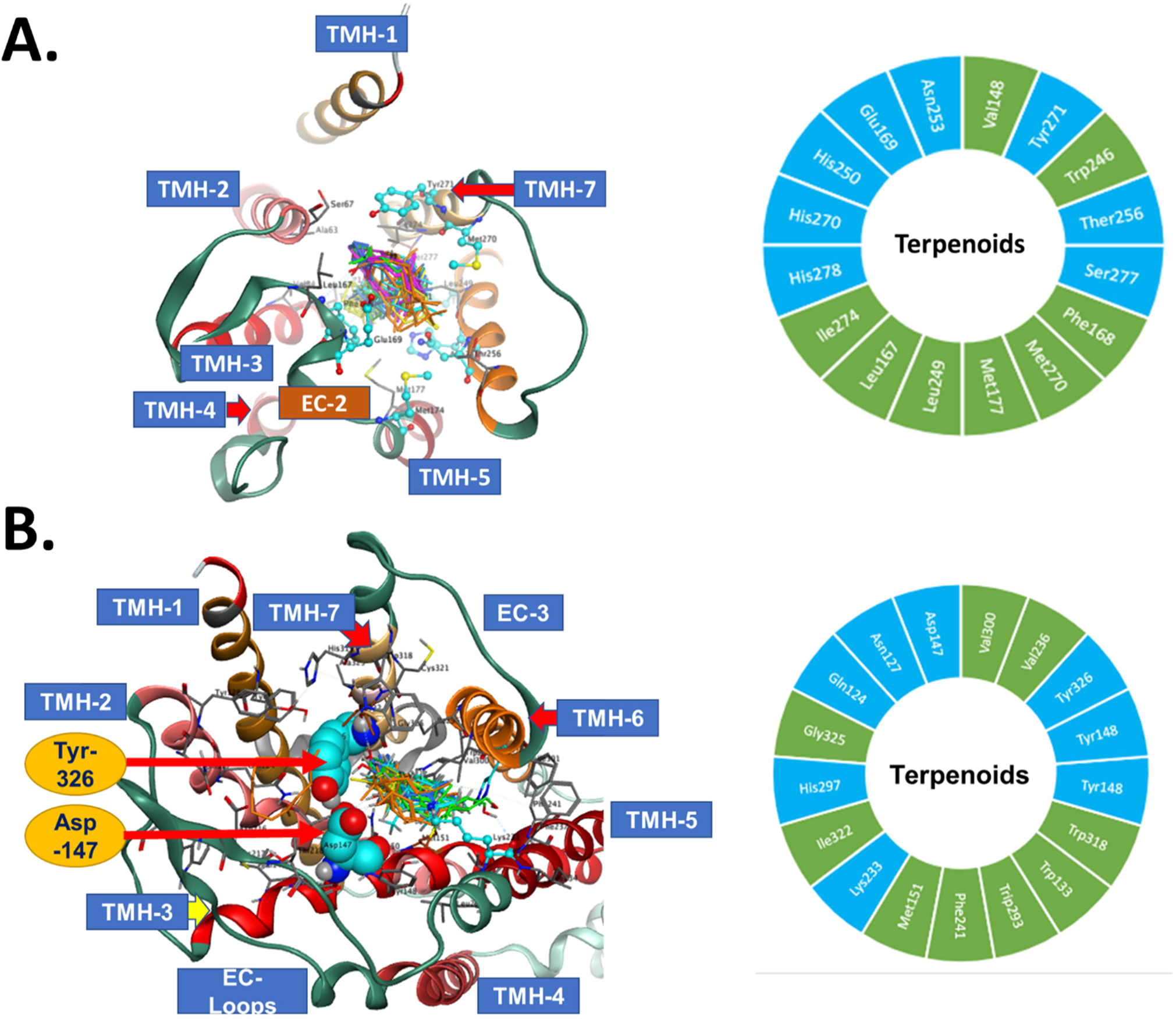
Molecular modeling of terpene binding with the A_2A_R and the mu opioid receptor (MOR). The 5 highest ranking poses of each terpene were docked to the A_2A_R and the MOR as described in the Methods. TMH = transmembrane helix; EC = extracellular domain. Docked illustrations are shown in the top-down view of the orthosteric binding pocket. **A)** *Left* – Docking poses of the terpenes in the A_2A_R binding pocket, along with noted interacting residues. *Right* – The 16 high confidence interacting residues for the terpenes are shown. Green = hydrophobic residues, Cyan = hydrophilic residues. **B)** *Left* – Docking poses of the terpenes in the MOR binding pocket, along with noted interacting residues. Tyr326 and Asp147 are particularly noted, since the terpenes lack key bonds with these residues, which may explain their lack of activity at the MOR. *Right* – The 17 high confidence interacting residues for the terpenes are shown. Green = hydrophobic residues, Cyan = hydrophilic residues.

For the MOR, 17 interacting amino acids at the binding cavity showed high binding energy and short interacting distances (**Figure 11B**). The distribution of amino acids was 9 hydrophobic and 8 hydrophilic amino acids. The difference in polarity at the binding sites of the A_2A_R and MOR is relatively small even though their natural ligands adenosine and morphine are more hydrophilic than lipophilic. Several unique amino acids of the MOR cavity interacted with the terpenes. Asp147(TMH-3) colored cyan and rendered spacefilling as well as Tyr326(TMH-7) are labeled in **Figure 11B**. Both residues are essential in the recognition of active opioid ligands [29]. Active opioids form ionic interactions with the side chain of Asp147(TMH-3) and the interaction is stabilized by Tyr326(TMH-7) [29]. In the docking of terpenes into MOR Asp147(TMH-3) and Tyr326(TMH-7) were observed interacting with geraniol and linalool by forming hydrogen bonds but no ionic interactions since none of the terpenes has an amine group in their structures. This lack of ionic interaction at these key residues may explain why our pharmacological data shows A_2A_R activation but no MOR activation, despite similar docking energies in **Table 1**. Overall, these *in vitro* and *in silico* studies suggested that all terpenes can directly activate the A_2A_R in order to cause the pain relief in CIPN observed above, and suggested a molecular hypothesis for this interaction.

### Terpenes May Have an Anti-Inflammatory Mechanism of Action

In addition to our A_2A_R mechanism identified above, literature reports have suggested various terpenes can have anti-inflammatory activity [12]. We thus hypothesized that these terpenes could produce antinociception in our acute LPS-induced pain model above by an anti-inflammatory mechanism. We first established proof-of-principle *in vitro* by testing the ability of the terpenes to reduce LPS stimulation of an NFκB- luciferase reporter in mouse BV-2 microglial cells. Notably, *all* terpenes tested produced significant anti-inflammatory activity in this *in vitro* model (**Figure S12A**). WIN55,212, linalool, and β-caryophyllene produced ∼50% inhibition of inflammatory activity, while geraniol, β-pinene, and α-humulene produced a complete blockade of LPS stimulation (**Figure S12A**). In order to rule out confounding effects on cell proliferation and/or cell death, we also performed a cell viability assay under the same treatment conditions. While β-pinene and α-humulene did cause a ∼30-50% decrease in cell viability, the other terpenes had little to no impact on cell viability compared to vehicle control (**Figure S12B**). These results thus suggest that most terpenes have a potential, genuine anti-inflammatory activity vs. LPS stimulation.

Following this, we next tested for changes in cytokine expression in the skin of mice treated with LPS and terpene in our acute inflammatory pain studies. We did find that geraniol, α-humulene, and β-pinene (p = 0.053) caused an elevation in levels of Interleukin-10 (IL-10), an anti-inflammatory cytokine (**Figure S13**). However, no terpene caused any changes in Tumor Necrosis Factor-α expression, and geraniol elevated levels of IL-6, while other terpenes had no impact on IL-6 expression (**Figure S13**). We thus could not observe a clear and consistent impact on cytokine levels across the different terpenes in our LPS pain model. We conclude that terpenes may have anti-inflammatory activity during *in vivo* pain models, but this cannot be conclusively demonstrated from our observations.

## Discussion

We report here that select terpenes from *Cannabis sativa* are efficacious in relieving neuropathic and inflammatory pain in mice. They consistently lack reward liability, enhance opioid pain relief, and produce equivalent antinociceptive tolerance to morphine. Our observations further suggest that terpenes produce pain relief in neuropathic pain via agonist activation of A_2A_ receptors in the spinal cord. Our model for these findings is summarized in **Figure 12**.

**Figure 12:**
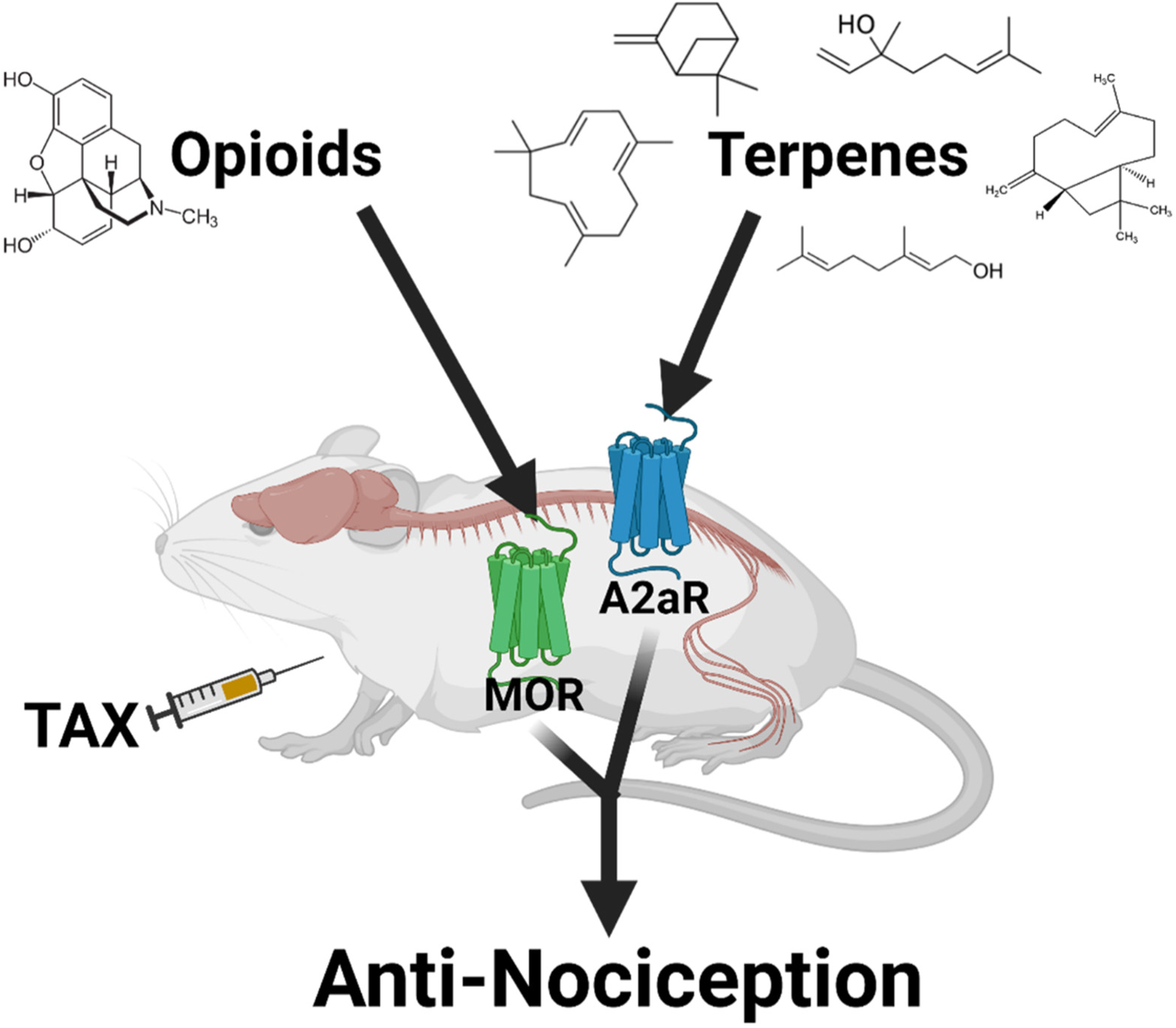
Model of terpene antinociception in CIPN. Under the condition of paclitaxel (TAX) treatment, terpene injection results in activation of the A_2A_R in the spinal cord, leading to antinociception. Separately, opioid injection can activate the MOR at an undefined site which enhances terpene pain relief.

This study was conceptualized in part to evaluate the translational potential of terpenes as analgesics in chronic pain. Besides the solo efficacy we show in CIPN and inflammatory pain, our data also showed that all terpenes enhanced the pain relief properties of morphine in CIPN. Analgesic additivity/synergy is a key translational benefit, which permits a dose-reduction strategy that maximizes pain relief while minimizing side effects. This has been shown for a number of drug combinations, including cannabinoid/opioid [27], adrenergic/opioid [30, 31], and different combinations of opioid subtype ligands (i.e. kappa and delta) [32]. The mechanism for our observed enhancement is not clear, although one report demonstrated that opioid treatment caused the release of adenosine from spinal synaptosomes, which could be a potential explanation [33]. Whatever the mechanism, this data suggests that low dose terpene/opioid combination therapy could provide efficacious pain relief with reduced side effects.

One unusual feature of this observed enhancement was that while terpenes enhanced opioid pain relief, they did not enhance cannabinoid pain relief, in sharp contrast to our earlier work which showed terpene/cannabinoid enhancement in tail flick pain [13]. One key difference is that our earlier work showed that terpene pain relief in tail flick pain was mediated by the CBR1, while here we show CIPN pain relief is mediated by the A_2A_R. Thus, perhaps in tail flick pain both ligands increase activation of their shared receptor target CBR1, which does not apply in CIPN pain. Alternatively, perhaps the CBR1 and A_2A_R systems do not interact in circuits relevant to CIPN pain, while opioid and A_2A_R systems do (as mentioned above [33]). Whatever the case, future work will need to disentangle how terpenes evoke different interacting receptor systems in different pain states. Another key translational feature we tested was reward liability by CPP; all terpenes lacked reward liability, while a few did show aversion liability. Few studies have tested the affective properties of terpenes, and those that have mostly did so in terms of their effects on other drugs of abuse [23–26, 34]. Promisingly, all those studies did show that the terpenes tested reduced place preference and self-administration of drugs of abuse like cocaine, although β-caryophyllene was the only terpene tested that was also tested in our studies above. Taken together, our findings thus suggest that terpenes alone have no abuse potential, and could reduce the abuse potential of other drugs like opioids (while enhancing their pain relief). In support of this hypothesis, a number of studies have shown that A_2A_R agonists can reduce place preference and self-administration of drugs of abuse [35]. This suggests that our terpenes could act through the A_2A_R expressed in the striatum to reduce the reward liability of drugs like morphine.

To our knowledge, translational features like analgesic tolerance have never been tested for terpenes. Here we show that the terpenes have comparable or slightly slower tolerance in CIPN than morphine. We also found that lower doses of terpene had less initial efficacy but also less apparent tolerance development; this is also true for other analgesic drugs, including opioids [36]. Unfortunately, we could not find studies that directly measured analgesic tolerance in response to A_2A_R agonist treatment, so we have no basis for contextualizing terpene analgesic tolerance at the A_2A_R. Analgesic tolerance has been shown for related receptors, such as with a Adenosine A_1_ positive allosteric modulator in spinal nerve ligation neuropathic pain [37]; this provides some reason to believe that analgesic tolerance is normal at the closely related A_2A_R. Future work will need to explore the mechanisms of analgesic tolerance at the A_2A_R and with terpenes in greater detail as these ligands move towards clinical use.

Our last translational aspect of study was preliminary dosing route testing, in which we found that our terpenes had limited bioavailability by the oral and inhaled routes. As we develop terpenes for clinical use, we propose formulation efforts to improve their pharmacokinetics at these routes. Potential options include formulation into nanoparticles made of various excipients, including polymers (e.g., polyethylene glycol) and lipids [38–41]. Such nanoparticle approaches were shown to make even metabolically labile ligands like peptides both metabolically stable, sometimes for days or weeks, as well as blood-brain barrier penetrant. Another potential option we did not explore here was transdermal delivery. Transdermal approaches generally use hydrophobic components to aid skin penetration, and can also be combined with nanoparticle formulation [42–44]. This approach may be particularly promising for terpenes, as one study showed that combining terpinolene with nanoparticles enhanced skin penetration [45]. Such approaches may be employed clinically to overcome the limitations we found here with unformulated, raw terpenes.

Beyond translation, we also sought to identify a mechanism of action for the antinociceptive activity of our terpenes, finding that they all likely act as A_2A_R agonists in CIPN, most in the spinal cord. As mentioned above, few studies have elucidated molecular mechanisms for any terpene in pain [12]. No study has connected any terpene to the A_2A_R in pain; limonene was identified as an A_2A_R agonist with anti-inflammatory and anti-anxiety activity, but this mechanism was not tested in pain [46–48]. Our findings linking terpenes to the A_2A_R in pain are thus novel. The A_2A_R itself has been conclusively linked to pain regulation, however, the findings are in some sense contradictory. A number of studies have clearly shown that intrathecal injection of A_2A_R agonists is antinociceptive in neuropathic pain models, providing precedent for terpenes activating A_2A_R in the spinal cord to be antinociceptive in CIPN [49, 50]. Work from this group suggested that A_2A_R agonists achieve pain relief in part by upregulating IL-10, which is noteworthy considering that we saw IL-10 upregulation in skin with some terpene treatments [51]. However, treatment with the A_2A_R antagonist caffeine was also acutely antinociceptive in neuropathic pain [52], and A_2A_R knockout mice showed a general hypoalgesic response [53]. The role of the A_2A_R in pain appears to be complex, with both pro-nociceptive and anti-nociceptive elements depending on context. In any case, the literature supports our observed mechanism of terpenes activating the A_2A_R to produce pain relief in CIPN.

Our work also suggested that the terpenes are A_2A_R agonists, as they evoked cAMP accumulation by the A_2A_R. However, the exact nature of that agonism is not clear. The terpenes did not compete with an orthosteric radioligand (similar to our results with the CBR1 [13]), suggesting that they might be allosteric agonists. However, our modeling studies suggested a mechanism for the terpenes to bind and activate the A_2A_R in the orthosteric binding pocket. One possibility is that the terpenes bind to a subcompartment of the orthosteric site that permits orthosteric activation but avoids radioligand competition; analogous findings have been described for cannabinoids that compete with ^3^H-WIN55,212 but not ^3^H-CP55,940 at the CB1R (discussed in [54]). Another possibility is that our terpenes bind the orthosteric site in a competing pose with ^3^H-ZM241385 but have unfavorable pharmacodynamic features (affinity, K_ON/OFF_, etc.) that prevents effective displacement of the radioligand. Or, of course, since molecular modeling is suggestive and hypothesis generating but not direct observation, our terpenes may be true allosteric agonists. Future work will need to determine the molecular pharmacology mechanisms behind terpene agonism of the A_2A_R, using tools and resources like a recent predictive mapping of A_2A_R allosteric sites [55].

Another interesting question is why our terpenes specifically activate the A_2A_R and not the CBR1 in CIPN, when we know from above and our previous work that the terpenes can activate both receptors [13] and that agonists of both receptors are anti-nociceptive in neuropathic pain (our WIN55,212 data above and [49–51, 56]). We also know that our terpenes can effectively engage the CBR1 *in vivo* since our previous work showed that terpene anti-nociception in tail flick pain was CBR1-mediated [13]. One possibility is that if our terpenes are true allosteric agonists, they may require co-activation of the target receptor with endogenous ligands (adenosine, endocannabinoids). In this case, there may be insufficient endocannabinoid tone in the spinal cord to lead to activation, while there could be sufficient adenosine (especially after evoked release by opioid treatment [33]). In support of this hypothesis, a recent study found decreased levels of anandamide and 2-arachidonoylglycerol in the spinal cords of rats with CIPN [57]. Future work will need to disentangle these mechanisms of selective receptor engagement in different pain states.

Overall, our observations support the translational utility of terpenes as potential treatments for neuropathic pain, and have identified a novel A_2A_R-mediated mechanism of action in spinal cord. Further work will be needed to overcome the translational hurdles identified, such as limited oral/inhaled bioavailability, and to further explore the antinociceptive mechanisms of action of these ligands. These ligands hold strong potential as non-opioid, non-cannabinoid analgesics for difficult to treat pain conditions like neuropathy, that hold promise to overcome the limitations of current treatments.

## Acknowledgments

This work was supported by NIH R01AT011517 and institutional funds from the University of Arizona to JMS. We would like to acknowledge Drs. Tally Largent-Milnes and Todd Vanderah of the University of Arizona Comprehensive Pain and Addiction Center for the use of their behavioral equipment. JMS is an equity holder in *Teleport Pharmaceuticals, LLC* and *Botanical Results, LLC*, a local cannabidiol company; no company products or interests were tested in this study. The authors have no other relevant conflicts of interest to declare.

## Data Availability

All data required to interpret these findings is contained within the manuscript or supplementary data. Raw data is available upon request to the corresponding author.

## Author Contributions

AMS participated in project design, performed experiments, analyzed data, and co-wrote the manuscript. AK participated in project design, performed experiments, and analyzed the data. TB, RJH, AP, BL, ZG, MG-R, CAS, KC, TLA performed experiments and analyzed their data. FAA performed independent modeling experiments and analyzed their data. KAJ participated in project design and supervised ZG in the performance of their work. JMS participated in project design, analyzed data, supervised AMS, AK, TB, RJH, AP, BL, MG-R, CAS, KC, and TLA in the performance of their work, secured funding, and co-wrote the manuscript. All authors had editorial input into the manuscript.

## References

1. Yong, R.J., P.M. Mullins, and N. Bhattacharyya, Prevalence of chronic pain among adults in the United States. Pain, 2022. 163(2): p. e328–e332.

2. Yang, J., et al., The Modified WHO Analgesic Ladder: Is It Appropriate for Chronic Non-Cancer Pain? J Pain Res, 2020. 13: p. 411–417.

3. Finnerup, N.B., et al., Pharmacotherapy for neuropathic pain in adults: a systematic review and meta-analysis. Lancet Neurol, 2015. 14(2): p. 162–73.

4. Derry, S., et al., Fentanyl for neuropathic pain in adults. Cochrane Database Syst Rev, 2016. 10: p. CD011605.

5. Grace, P.M., et al., Protraction of neuropathic pain by morphine is mediated by spinal damage associated molecular patterns (DAMPs) in male rats. Brain Behav Immun, 2017.

6. Hill, K.P., Medical Marijuana for Treatment of Chronic Pain and Other Medical and Psychiatric Problems: A Clinical Review. JAMA, 2015. 313(24): p. 2474–83.

7. Mahabir, V.K., et al., Medical cannabis use in the United States: a retrospective database study. J Cannabis Res, 2020. 2(1): p. 32.

8. Uberall, M.A., A Review of Scientific Evidence for THC:CBD Oromucosal Spray (Nabiximols) in the Management of Chronic Pain. J Pain Res, 2020. 13: p. 399–410.

9. Rabgay, K., et al., The effects of cannabis, cannabinoids, and their administration routes on pain control efficacy and safety: A systematic review and network meta-analysis. J Am Pharm Assoc (2003), 2020. 60(1): p. 225–234 e6.

10. Allen, K.D., et al., Genomic characterization of the complete terpene synthase gene family from Cannabis sativa. PLoS One, 2019. 14(9): p. e0222363.

11. Mudge, E.M., P.N. Brown, and S.J. Murch, The Terroir of Cannabis: Terpene Metabolomics as a Tool to Understand Cannabis sativa Selections. Planta Medica, 2019. 85(09/10): p. 781–796.

12. Liktor-Busa, E., et al., Analgesic Potential of Terpenes Derived from Cannabis sativa, in Pharmacological Reviews. 2021, American Society for Pharmacology and Experimental Therapeutics. p. 1269–1297.

13. LaVigne, J.E., et al., Cannabis sativa terpenes are cannabimimetic and selectively enhance cannabinoid activity. Sci Rep, 2021. 11(1): p. 8232.

14. Stine, C., et al., Heat shock protein 90 inhibitors block the antinociceptive effects of opioids in mouse chemotherapy-induced neuropathy and cancer bone pain models. Pain, 2020. 161(8): p. 1798–1807.

15. Chaplan, S.R., et al., Quantitative assessment of tactile allodynia in the rat paw. J Neurosci Methods, 1994. 53(1): p. 55–63.

16. Duron, D.I., F. Hanak, and J.M. Streicher, Daily intermittent fasting in mice enhances morphine-induced anti-nociception while mitigating reward, tolerance, and constipation. Pain, 2020.

17. Lei, W., et al., The Alpha Isoform of Heat Shock Protein 90 and the Co-chaperones p23 and Cdc37 Promote Opioid Anti-nociception in the Brain. Front Mol Neurosci, 2019. 12: p. 294.

18. Duron, D.I., et al., Inhibition of Hsp90 in the spinal cord enhances the antinociceptive effects of morphine by activating an ERK-RSK pathway. Sci Signal, 2020. 13(630).

19. Suresh, R.R., et al., Selective A(3) Adenosine Receptor Antagonist Radioligand for Human and Rodent Species. ACS Med Chem Lett, 2022. 13(4): p. 623–631.

20. Mossine, V.V., et al., piggyBac transposon plus insulators overcome epigenetic silencing to provide for stable signaling pathway reporter cell lines. PLoS One, 2013. 8(12): p. e85494.

21. Lei, W., et al., A Novel Mu-Delta Opioid Agonist Demonstrates Enhanced Efficacy with Reduced Tolerance and Dependence in Mouse Neuropathic Pain Models. J Pain, 2019.

22. Lei, W., et al., Heat shock protein 90 (Hsp90) promotes opioid-induced anti-nociception by an ERK Mitogen Activated Protein Kinase (MAPK) mechanism in mouse brain. J Biol Chem, 2017.

23. Galaj, E., et al., Beta-caryophyllene inhibits cocaine addiction-related behavior by activation of PPARalpha and PPARgamma: repurposing a FDA-approved food additive for cocaine use disorder. Neuropsychopharmacology, 2021. 46(4): p. 860–870.

24. He, X.H., et al., beta-caryophyllene, an FDA-Approved Food Additive, Inhibits Methamphetamine-Taking and Methamphetamine-Seeking Behaviors Possibly via CB2 and Non-CB2 Receptor Mechanisms. Front Pharmacol, 2021. 12: p. 722476.

25. Asth, L., et al., Effects of beta -caryophyllene, A Dietary Cannabinoid, in Animal Models of Drug Addiction. Curr Neuropharmacol, 2023. 21(2): p. 213–218.

26. Gu, S.M., et al., Limonene Inhibits Methamphetamine-Induced Sensitizations via the Regulation of Dopamine Receptor Supersensitivity. Biomol Ther (Seoul), 2019. 27(4): p. 357–362.

27. Grenald, S.A., et al., Synergistic attenuation of chronic pain using mu opioid and cannabinoid receptor 2 agonists. Neuropharmacology, 2017. 116: p. 59–70.

28. Garcia-Nafria, J., et al., Cryo-EM structure of the adenosine A(2A) receptor coupled to an engineered heterotrimeric G protein. Elife, 2018. 7.

29. Perlikowska, R., et al., Pharmacological properties of novel cyclic pentapeptides with micro-opioid receptor agonist activity. Med Chem, 2014. 10(2): p. 154–61.

30. Stone, L.S., et al., Morphine and clonidine combination therapy improves therapeutic window in mice: synergy in antinociceptive but not in sedative or cardiovascular effects. PLoS One, 2014. 9(10): p. e109903.

31. Chabot-Dore, A.J., M. Millecamps, and L.S. Stone, The delta-opioid receptor is sufficient, but not necessary, for spinal opioid-adrenergic analgesic synergy. J Pharmacol Exp Ther, 2013. 347(3): p. 773–80.

32. Sutters, K.A., et al., Analgesic synergy and improved motor function produced by combinations of mu- delta- and mu-kappa-opioids. Brain Res, 1990. 530(2): p. 290–4.

33. Cahill, C.M., T.D. White, and J. Sawynok, Synergy between mu/delta-opioid receptors mediates adenosine release from spinal cord synaptosomes. Eur J Pharmacol, 1996. 298(1): p. 45–9.

34. Pourtaqi, N., et al., Effect of linalool on the acquisition and reinstatement of morphine-induced conditioned place preference in mice. Avicenna J Phytomed, 2017. 7(3): p. 242–249.

35. Wydra, K., et al., Adenosine A(2A)Receptors in Substance Use Disorders: A Focus on Cocaine. Cells, 2020. 9(6).

36. Yoburn, B.C., B. Billings, and A. Duttaroy, Opioid receptor regulation in mice. J Pharmacol Exp Ther, 1993. 265(1): p. 314–20.

37. Li, X., et al., Repeated dosing with oral allosteric modulator of adenosine A1 receptor produces tolerance in rats with neuropathic pain. Anesthesiology, 2004. 100(4): p. 956–61.

38. Rhee, Y.S., et al., Sustained-release delivery of octreotide from biodegradable polymeric microspheres. AAPS PharmSciTech, 2011. 12(4): p. 1293–301.

39. Popov, M., et al., Delivery of analgesic peptides to the brain by nano-sized bolaamphiphilic vesicles made of monolayer membranes. Eur J Pharm Biopharm, 2013. 85(3 Pt A): p. 381–9.

40. Lindqvist, A., et al., Enhanced brain delivery of the opioid peptide DAMGO in glutathione pegylated liposomes: a microdialysis study. Mol Pharm, 2013. 10(5): p. 1533–41.

41. Patel, M., E.B. Souto, and K.K. Singh, Advances in brain drug targeting and delivery: limitations and challenges of solid lipid nanoparticles. Expert Opin Drug Deliv, 2013. 10(7): p. 889–905.

42. Giacoppo, S., et al., A new formulation of cannabidiol in cream shows therapeutic effects in a mouse model of experimental autoimmune encephalomyelitis. Daru, 2015. 23: p. 48.

43. Nitecka-Buchta, A., et al., Myorelaxant Effect of Transdermal Cannabidiol Application in Patients with TMD: A Randomized, Double-Blind Trial. J Clin Med, 2019. 8(11).

44. Nastiti, C., et al., Novel Nanocarriers for Targeted Topical Skin Delivery of the Antioxidant Resveratrol. Pharmaceutics, 2020. 12(2).

45. Kahraman, E., et al., The combination of nanomicelles with terpenes for enhancement of skin drug delivery. Int J Pharm, 2018. 551(1-2): p. 133–140.

46. Park, H.M., et al., Limonene, a natural cyclic terpene, is an agonistic ligand for adenosine A(2A) receptors. Biochem Biophys Res Commun, 2011. 404(1): p. 345–8.

47. Patel, M., et al., Limonene-induced activation of A(2A) adenosine receptors reduces airway inflammation and reactivity in a mouse model of asthma. Purinergic Signal, 2020. 16(3): p. 415–426.

48. Song, Y., et al., Limonene has anti-anxiety activity via adenosine A2A receptor-mediated regulation of dopaminergic and GABAergic neuronal function in the striatum. Phytomedicine, 2021. 83: p. 153474.

49. Kwilasz, A.J., et al., Sustained reversal of central neuropathic pain induced by a single intrathecal injection of adenosine A2A receptor agonists. Brain Behav Immun, 2018. 69: p. 470–479.

50. Loram, L.C., et al., Intrathecal injection of adenosine 2A receptor agonists reversed neuropathic allodynia through protein kinase (PK)A/PKC signaling. Brain Behav Immun, 2013. 33: p. 112–22.

51. Loram, L.C., et al., Enduring reversal of neuropathic pain by a single intrathecal injection of adenosine 2A receptor agonists: a novel therapy for neuropathic pain. J Neurosci, 2009. 29(44): p. 14015–25.

52. Wu, W.P., et al., Effect of acute and chronic administration of caffeine on pain-like behaviors in rats with partial sciatic nerve injury. Neurosci Lett, 2006. 402(1-2): p. 164–6.

53. Hussey, M.J., et al., Reduced response to the formalin test and lowered spinal NMDA glutamate receptor binding in adenosine A2A receptor knockout mice. Pain, 2007. 129(3): p. 287–294.

54. Franco, R., et al., The Binding Mode to Orthosteric Sites and/or Exosites Underlies the Therapeutic Potential of Drugs Targeting Cannabinoid CB(2) Receptors. Front Pharmacol, 2022. 13: p. 852631.

55. Caliman, A.D., Y. Miao, and J.A. McCammon, Mapping the allosteric sites of the A(2A) adenosine receptor. Chem Biol Drug Des, 2018. 91(1): p. 5–16.

56. King, K.M., et al., Single and combined effects of Delta(9) -tetrahydrocannabinol and cannabidiol in a mouse model of chemotherapy-induced neuropathic pain. Br J Pharmacol, 2017. 174(17): p. 2832–2841.

57. Noya-Riobo, M.V., et al., Changes in the expression of endocannabinoid system components in an experimental model of chemotherapy-induced peripheral neuropathic pain: Evaluation of sex-related differences. Exp Neurol, 2023. 359: p. 114232.

