## Supplementary Data for "Terpenes from *Cannabis sativa* Induce Antinociception in Mouse Chronic Neuropathic Pain via Activation of Spinal Cord Adenosine A_2A_ Receptors"

**Running Title:** Terpenes Relieve Chronic Pain via the $A_{2A}$ Receptor

**Keywords:** Terpenes; Cannabis; Neuropathic Pain; Adenosine $A_{2A}$ Receptor; Analgesia

### Figures

#### Figure S1:

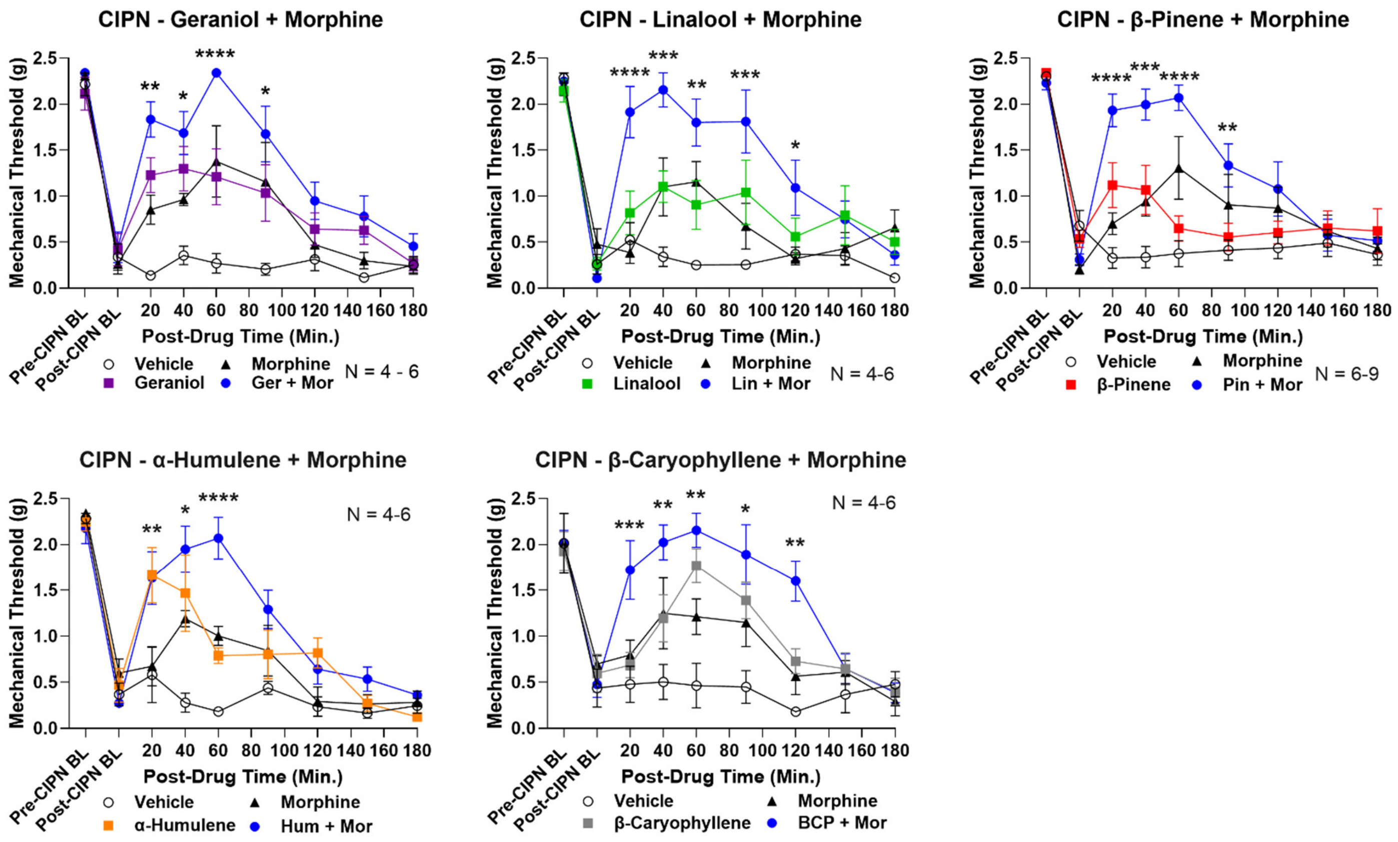


**Figure S1: Combination of low dose terpene and morphine enhances antinociception in CIPN.** Male and female CD-1 mice had CIPN induced and measured as described in the Methods. Data shown is the mean ± SEM, performed in 2-3 technical replicates for each experiment, with sample sizes noted in each graph. BL = baseline. Vehicle, morphine (3.2 mg/kg, SC), terpene (100 mg/kg, IP) or both combined were injected, with mechanical allodynia measured. *, **, ***, **** = $p < 0.05, 0.01, 0.001, 0.0001$ vs. same time point terpene or morphine alone group by RM 2 Way ANOVA with Dunnett's *post hoc* test. Terpene/morphine combination enhances pain relief over either alone.

**Figure S2:**

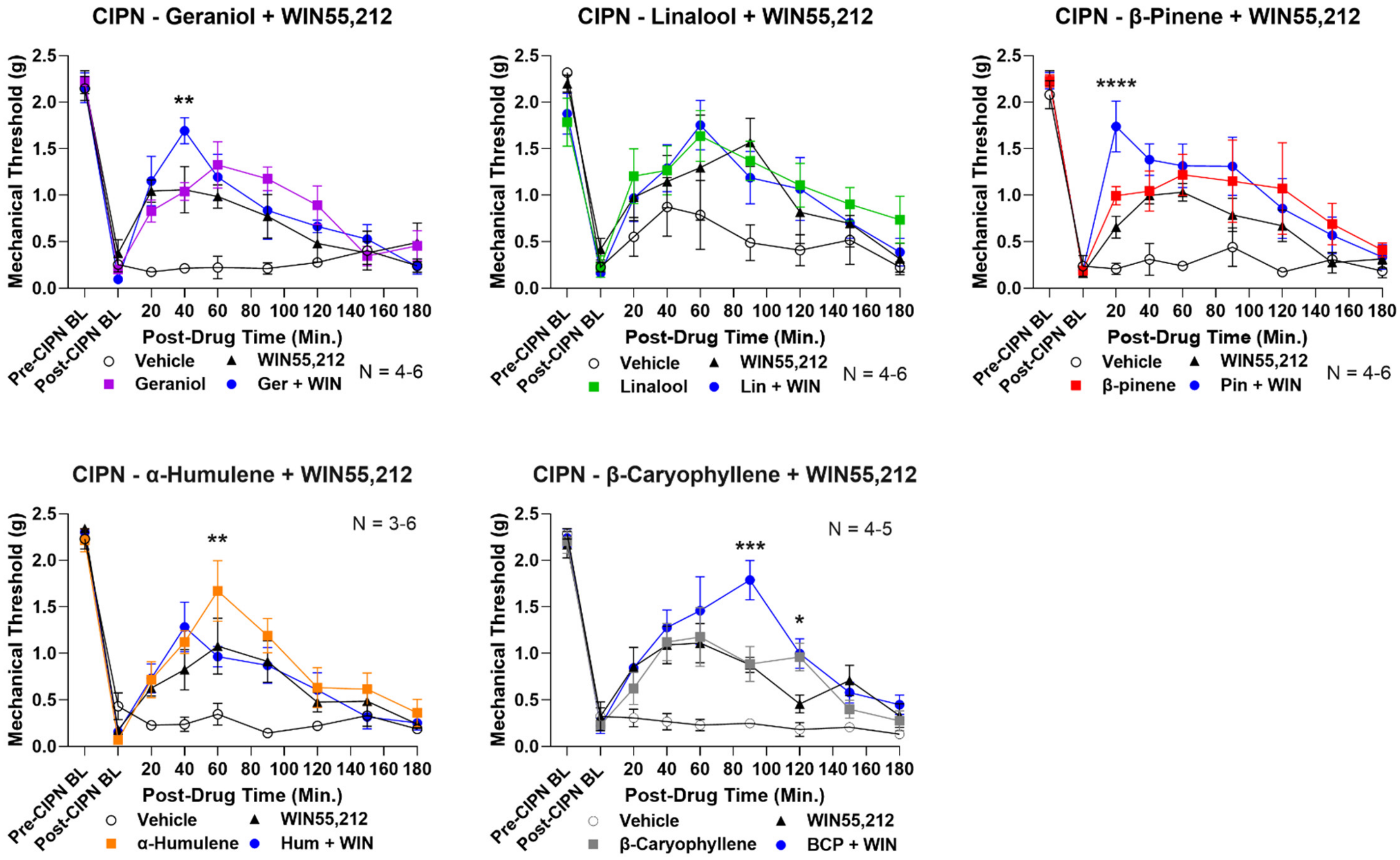


**Figure S2: Combination of low dose terpene and cannabinoid has little to no enhancement of pain relief.** Male and female CD-1 mice had CIPN induced and measured as described in the Methods. Data shown is the mean ± SEM, performed in 2-3 technical replicates for each experiment, with sample sizes noted in each graph. BL = baseline. Vehicle, WIN55,212 (1 mg/kg, IP), terpene (100 mg/kg, IP) or both combined were injected, with mechanical allodynia measured. *, **, ***, **** = $p < 0.05$, 0.01, 0.001, 0.0001 vs. same time point terpene or WIN55,212 alone group by RM 2 Way ANOVA with Dunnett's *post hoc* test. Terpene/WIN55,212 combination had little to no enhancing effect on pain relief in CIPN.

**Figure S3:**

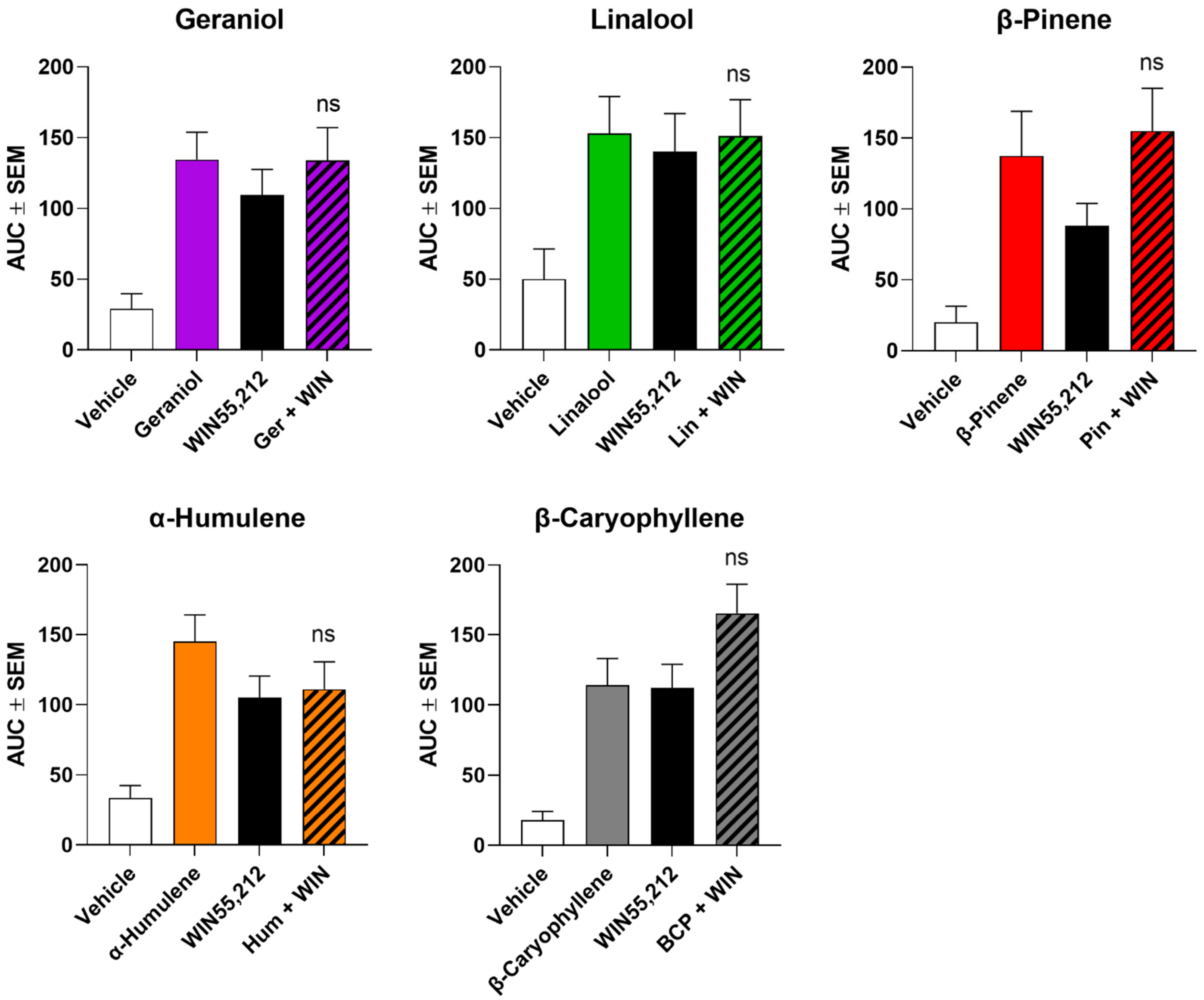


**Figure S3: AUC data from terpene/WIN55,212 combination experiments.** AUC data calculated from the time course curves in **Figure S2**. Terpene/WIN55,212 combination had no significant enhancement for any terpene ($p > 0.05$ by 1 Way ANOVA with Tukey's *post hoc* test).

**Figure S4:**

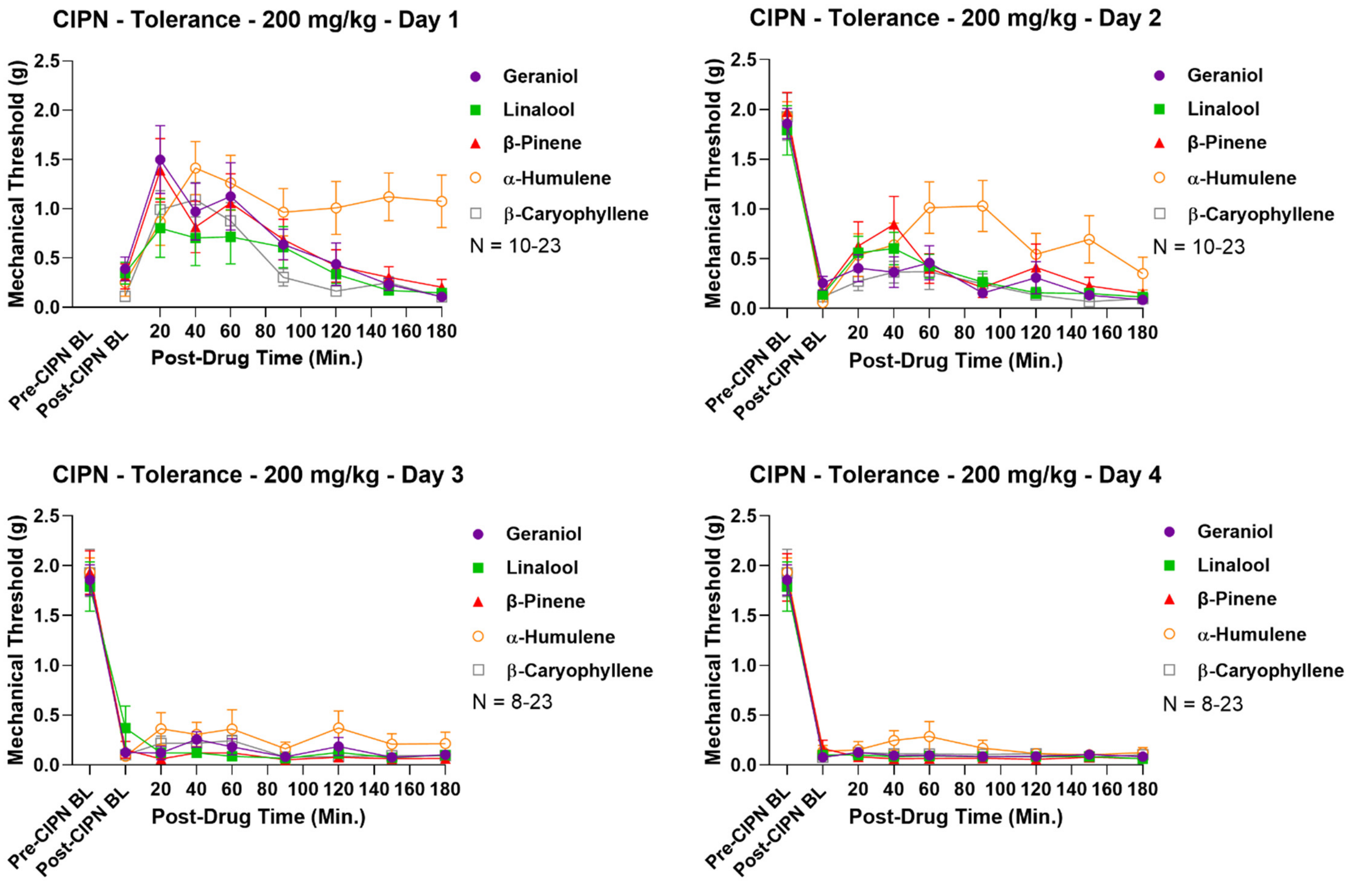


**Figure S4: Raw time course data for terpene tolerance testing at 200 mg/kg.** Male and female CD-1 mice had CIPN induced as in the Methods, with terpene (200 mg/kg, IP) injection twice daily over a 4-day protocol, with daily mechanical allodynia measurement after the morning injection. Data shown is the mean ± SEM, performed in 2-5 technical replicates for each experiment, with sample sizes noted in each graph. BL = baseline.

**Figure S5:**

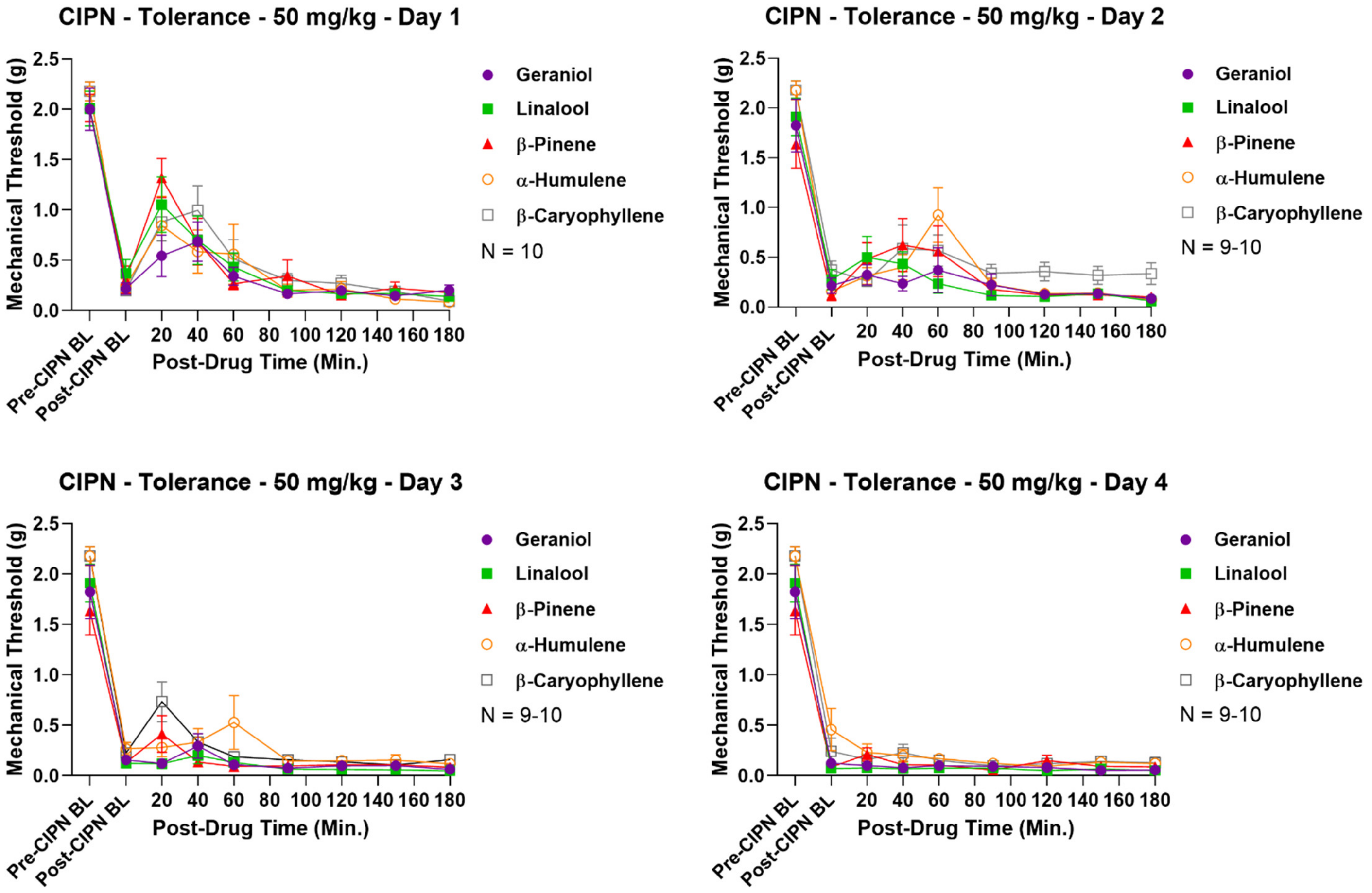


**Figure S5: Raw time course data for terpene tolerance testing at 50 mg/kg.** Male and female CD-1 mice had CIPN induced as in the Methods, with terpene (50 mg/kg, IP) injection twice daily over a 4-day protocol, with daily mechanical allodynia measurement after the morning injection. Data shown is the mean ± SEM, performed in 2 technical replicates for each experiment, with sample sizes noted in each graph. BL = baseline.

**Figure S6:**

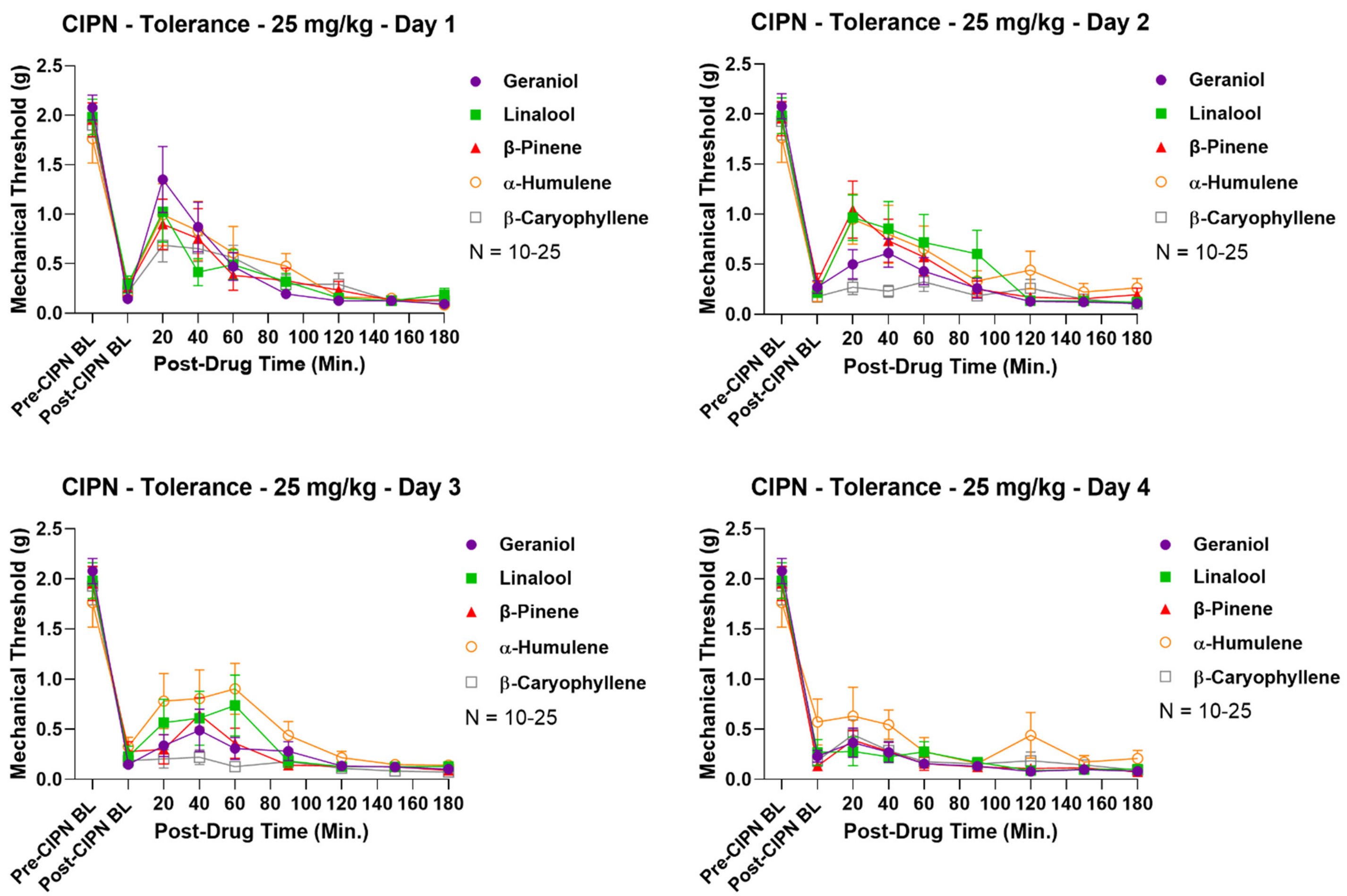


**Figure S6: Raw time course data for terpene tolerance testing at 25 mg/kg.** Male and female CD-1 mice had CIPN induced as in the Methods, with terpene (25 mg/kg, IP) injection twice daily over a 4-day protocol, with daily mechanical allodynia measurement after the morning injection. Data shown is the mean ± SEM, performed in 2-5 technical replicates for each experiment, with sample sizes noted in each graph. BL = baseline.

**Figure S7:**

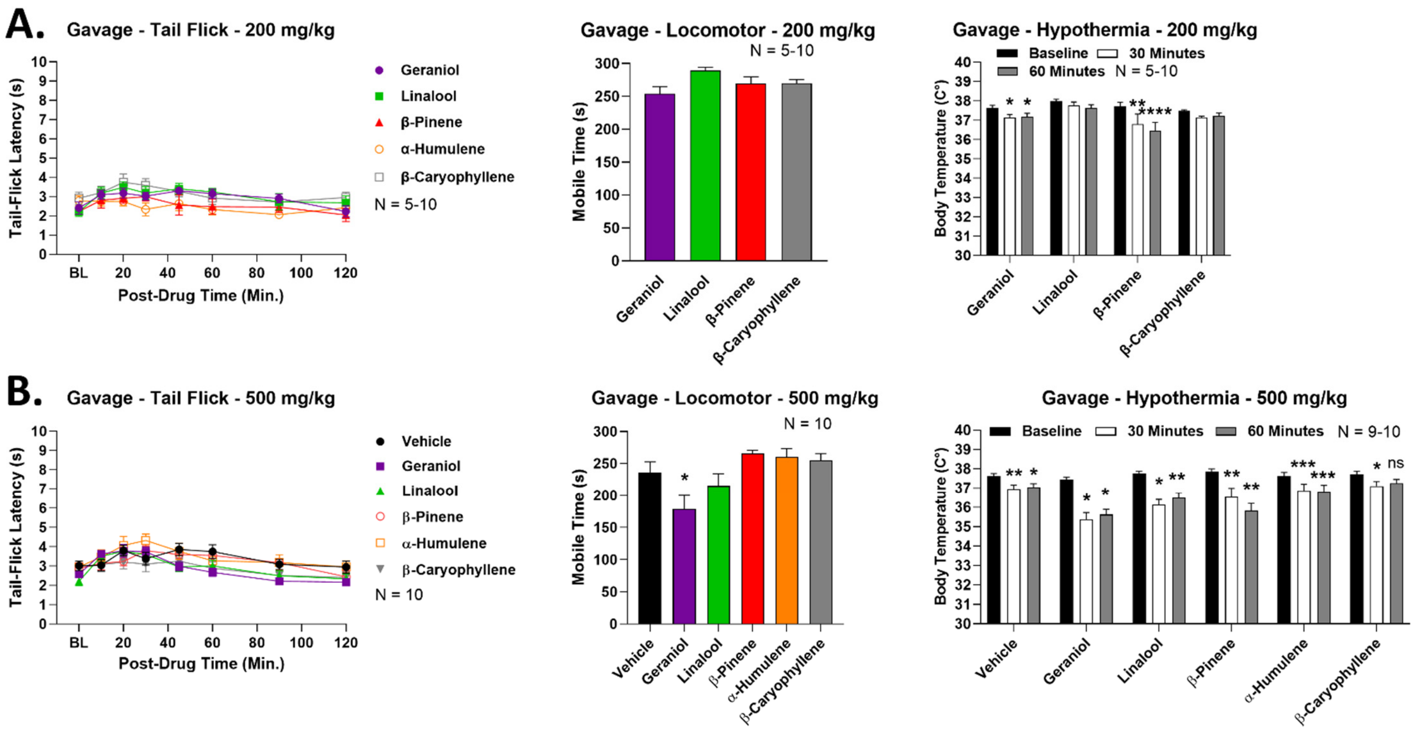


**Figure S7: Terpene bioavailability testing by the oral route.** Male and female CD-1 mice had terpene (200-500 mg/kg, PO) or vehicle delivered by oral gavage, followed by cannabinoid tetrad testing (see Methods). Data shown is the mean ± SEM performed over 1-2 technical replicates, with sample sizes noted in each graph. BL = baseline. **A)** 200 mg/kg dose data. *, **, **** = $p < 0.05, 0.01, 0.0001$ vs. same terpene baseline measurement by RM 2 Way ANOVA with Dunnett's *post hoc* test. Only hypothermia showed any significant effects, so full testing with all terpenes was not performed. **B)** 500 mg/kg dose data. Locomotor: * = $p < 0.05$ vs. vehicle group by 1 Way ANOVA with Dunnett's *post hoc* test. Hypothermia: *, **, *** = $p < 0.05, 0.01, 0.001$ vs. same treatment baseline measurement by RM 2 Way ANOVA with Dunnett's *post hoc* test. The fully powered testing was completed at 500 mg/kg, but only limited bioavailability was shown.

**Figure S8:**

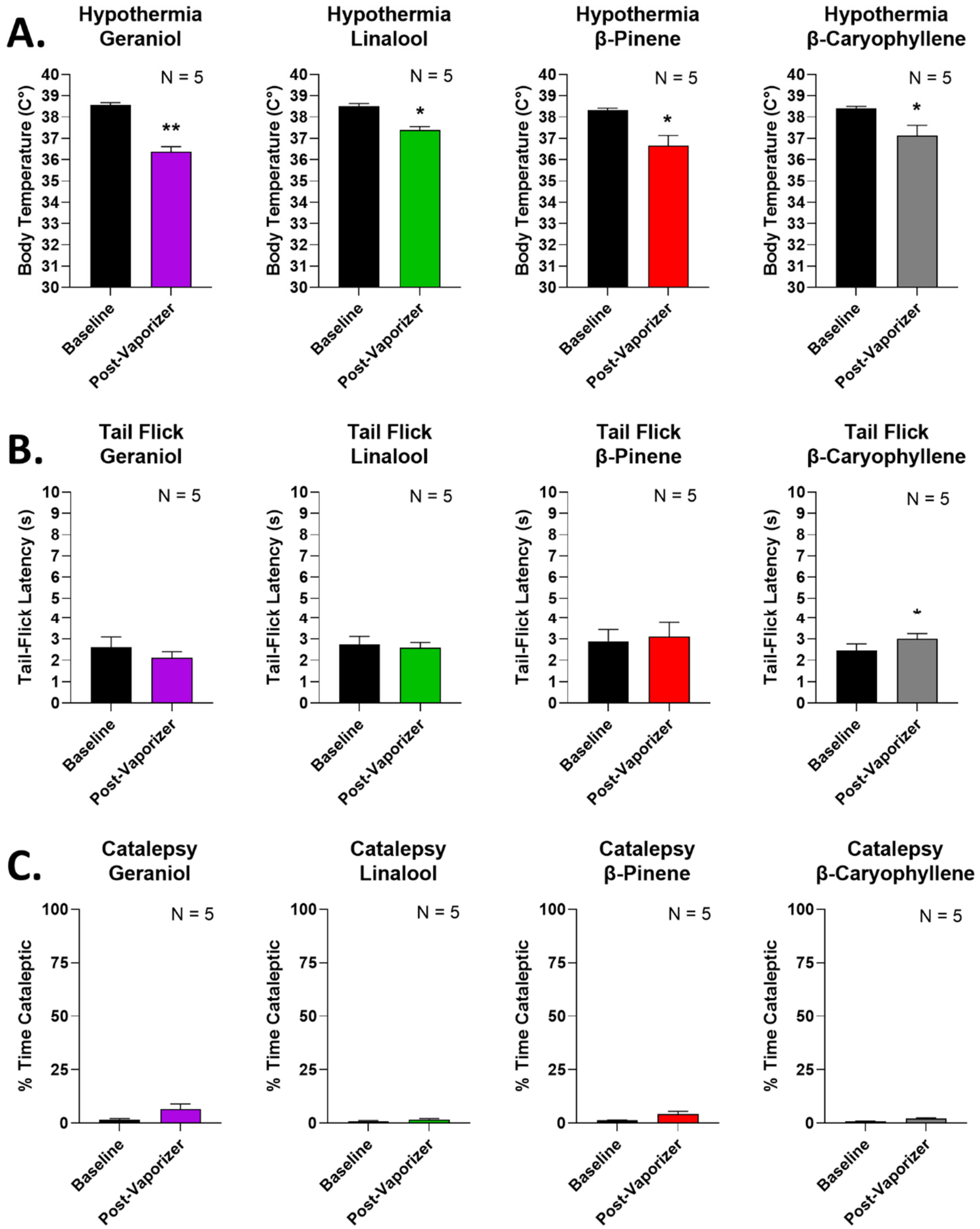


**Figure S8: Terpene bioavailability testing by the inhaled route.** Male and female CD-1 mice were exposed to pure vaporized terpene for 20 min followed by cannabinoid tetrad testing (see Methods). Data expressed as the mean ± SEM performed in 1 technical replicate, with sample sizes noted in each graph. **A)** Hypothermia measurement. *, ** = $p < 0.05$, 0.01 vs. baseline measurement by Paired 2-Tailed *t* Test. **B)** Tail flick response data. * = $p < 0.05$ vs. baseline measurement by Paired 2-Tailed *t* Test. **C)** Catalepsy data. No significant differences with treatment ($p > 0.05$). Since the response to inhaled terpene also showed limited bioavailability, we did not fully power this study with all terpenes.

**Figure S9:**

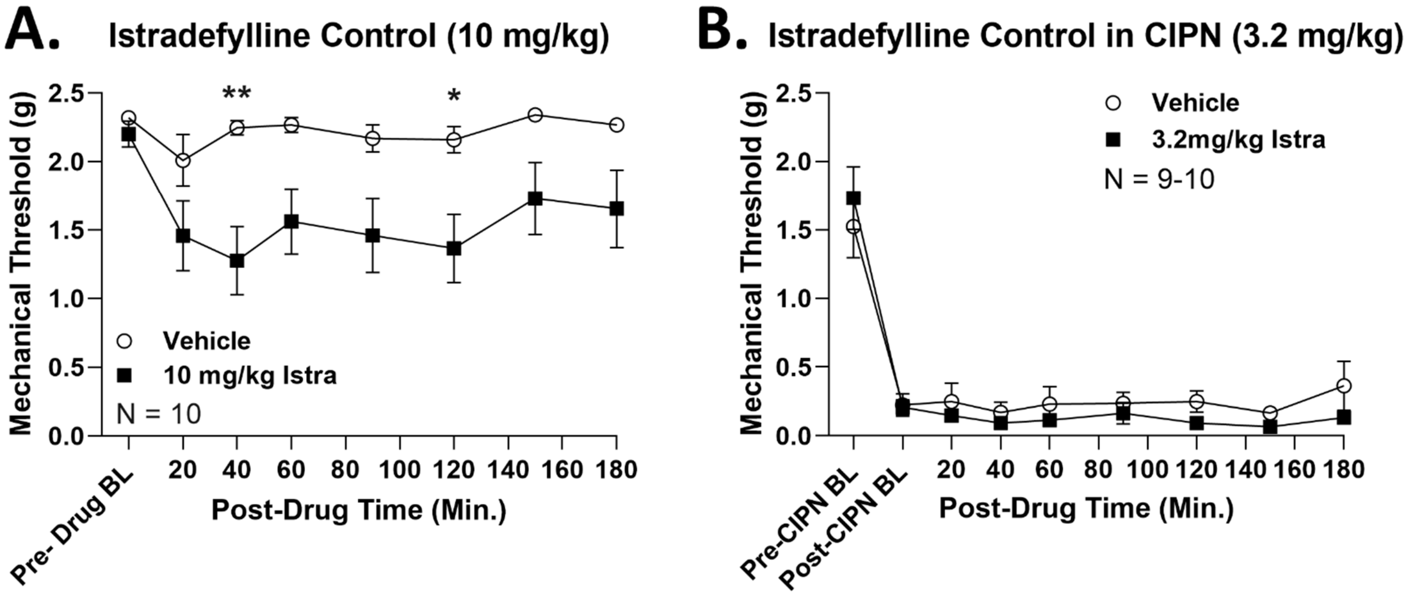


**Figure S9: Validation of istradefylline dose used in behavioral testing.** Male and female CD-1 mice used. Data represented as the mean ± SEM performed in 2 technical replicates, with sample sizes shown in each graph. BL = baseline. **A)** Naïve mice (no CIPN) injected with vehicle or istradefylline (10 mg/kg, IP) and mechanical thresholds measured. *, ** = $p < 0.05$, 0.01 vs. same time point istradefylline group by RM 2 Way ANOVA with Sidak's *post hoc* test. Istradefylline produced mechanical sensitivity in naïve mice, so this dose could not be used for behavioral testing. **B)** CIPN mice injected with vehicle or istradefylline (3.2 mg/kg, IP), and mechanical thresholds measured. Istradefylline had no effect on CIPN mechanical thresholds ($p > 0.05$), further validating this lower dose for behavioral testing.

**Figure S10:**

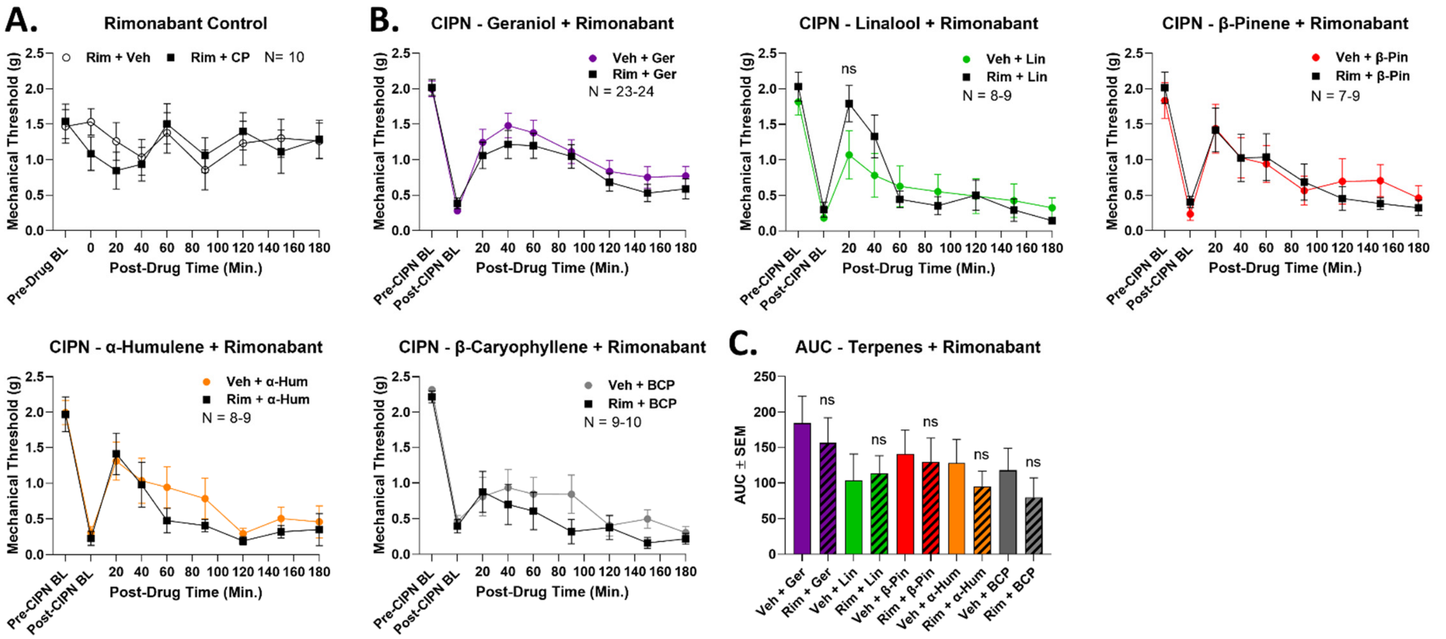


**Figure S10: Terpenes do not produce antinociception via the CBR1 in CIPN.** Male and female CD-1 mice had CIPN induced. On day 8, vehicle or rimonabant (10 mg/kg, IP) injected with a 10 min treatment, followed by terpene (200 mg/kg, IP) and mechanical threshold measurements over a 3 hr time course. Data represented as the mean ± SEM performed in 2-5 technical replicates, with sample sizes shown in each graph. BL = baseline. **A)** Rimonabant dose validation. Naïve mice (no CIPN) had vehicle or rimonabant (10 mg/kg, IP) injected with a 10 min treatment, followed by vehicle or CP55,940 (3.2 mg/kg, IP) and mechanical threshold measurement. Rimonabant had no impact on any baseline response, validating the drug dose ($p > 0.05$). **B)** Rimonabant and terpene injected in CIPN as described above. Rimonabant had no significant effect on any terpene response ($p > 0.05$). **C)** AUC data calculated from **B**. Rimonabant had no impact on any terpene response ($p > 0.05$).

**Figure S11:**

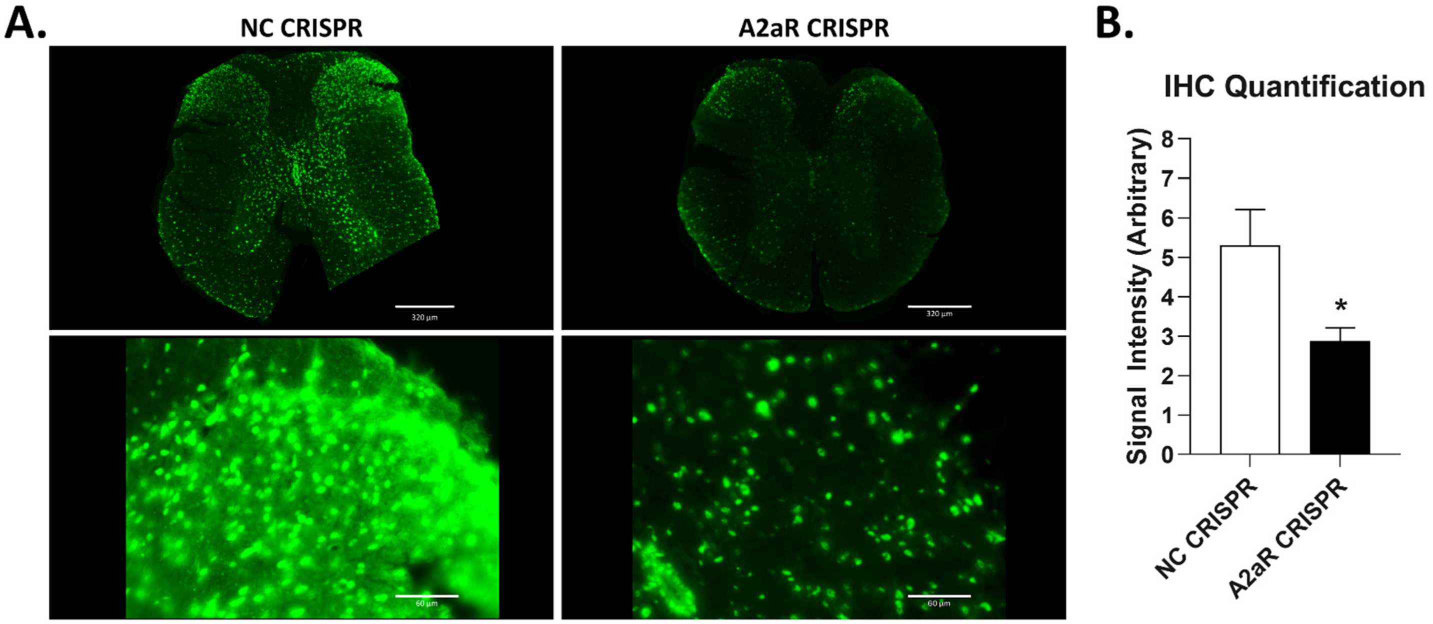


**Figure S11: Validation of CRISPR-mediated $A_{2A}R$ knockdown in spinal cord.** Male and female CD-1 mice had Negative Control (NC) or $A_{2A}R$-targeted CRISPR treatment as described in the Methods. The spinal cords were perfused and fixed, and stained for the $A_{2A}R$ (green). **A)** Representative images of whole spinal cord (*top*) and dorsal horn region (*bottom*) from both CRISPR treatments. There is an apparent knockdown of $A_{2A}R$ expression across the tissue. **B)** Quantitation of $A_{2A}R$ fluorescence from both conditions using ImageJ (see Methods). At least 2 sections per mouse were quantified, and those values averaged to produce a single value per mouse. N = 4 mice/group. * = $p < 0.05$ vs. NC group by Unpaired 2-Tailed *t* Test. $A_{2A}R$-targeted CRISPR significantly reduced the expression of the receptor in the spinal cord.

**Figure S12:**

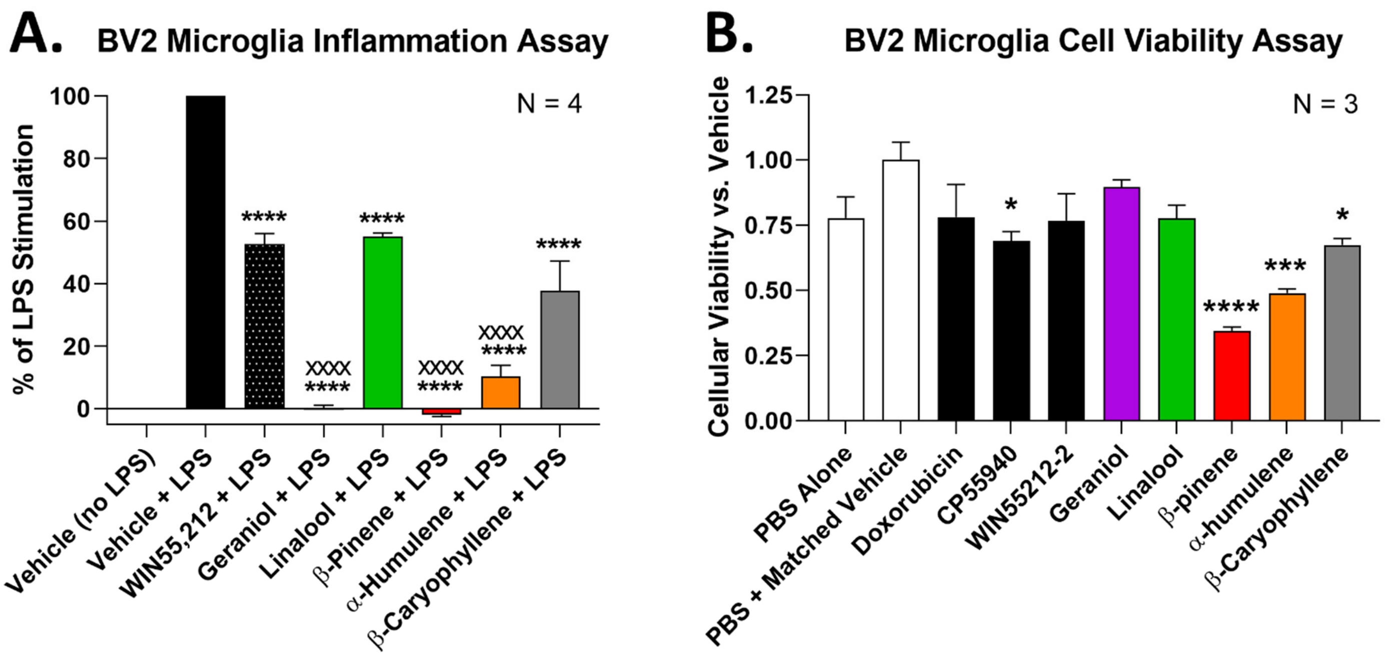


**Figure S12: Terpenes reduce inflammatory activation in BV2 microglial cells.** Data represents the mean ± SEM of 3-4 independent technical replicates. Sample sizes of independent experiments shown in each graph. **A)** BV2 cells with an NFκB-luciferase reporter used. Cells were serum starved (DMEM media alone) for 1 hour, pre-treated with 500 μM of individual terpene, 10 μM WIN55,212, or matched vehicle for 5 minutes, after which 1 μg/mL of LPS was added for 3 hrs at 37°C followed by luciferase measurement. **** = $p < 0.0001$ vs. Vehicle/LPS group; xxxx = $p < 0.0001$ vs. WIN55,212/LPS group; both by 1 Way ANOVA with Tukey's *post hoc* test. **B)** Resazurin cell viability assay performed to rule out confounding effects on proliferation/cell death. 500 μM terpene, 10 μM doxorubicin, 10 μM CP55,940, 10 μM WIN55,212, or matched vehicle (0.5% DMSO) were added to appropriate wells and incubated for 1 hour at 37°C. 0.075 mg/mL resazurin was added to each well and incubated for a further 2 hours at 37°C. *, ***, **** = $p < 0.05$, 0.001, 0.0001 vs. PBS/Vehicle group by 1 Way ANOVA with Tukey's *post hoc* test. Most terpenes had little to no effect on cell viability, validating the inflammatory assay in **A**.

**Figure S13:**

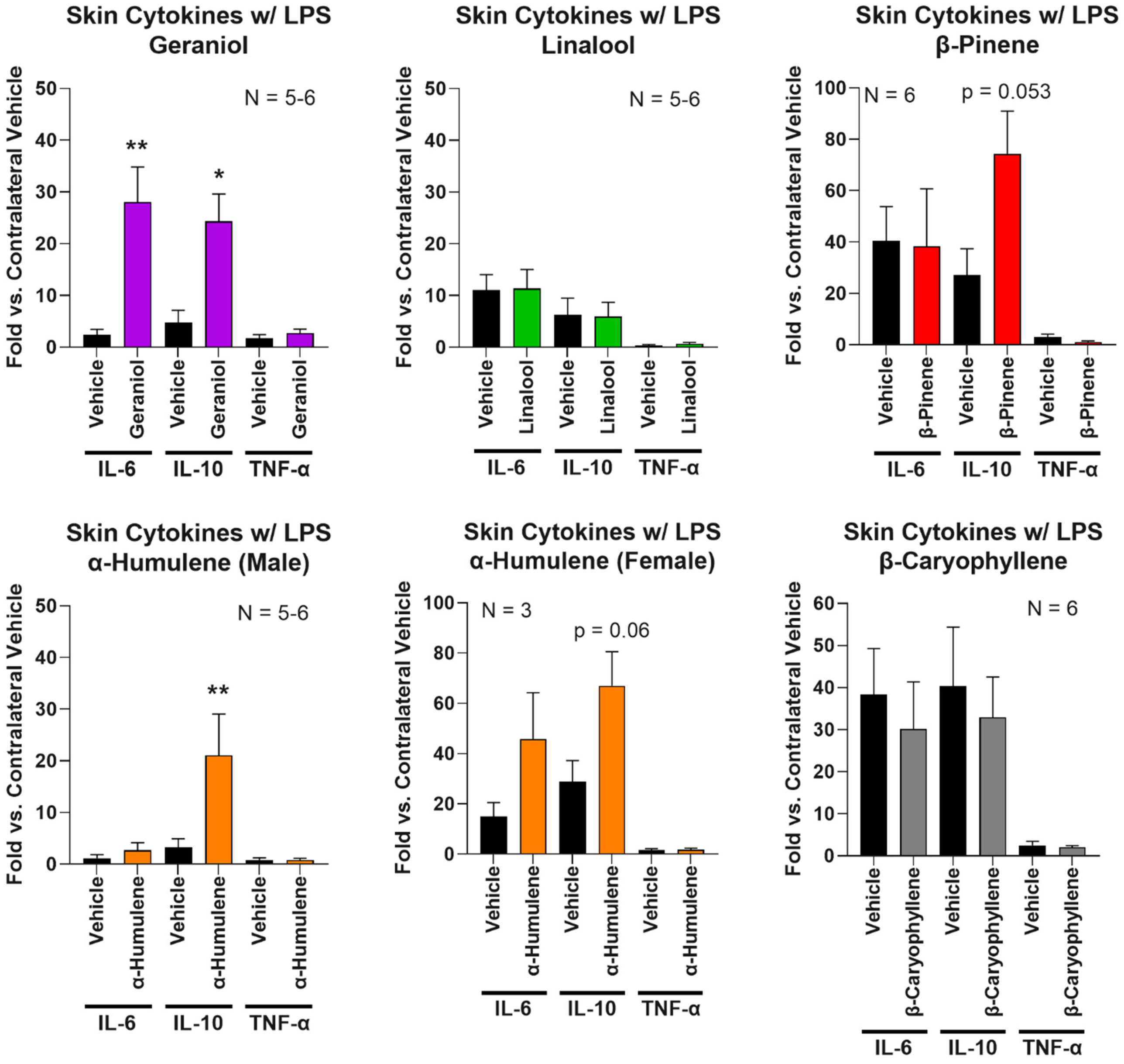


**Figure S13: Some terpenes may have an anti-inflammatory effect in acute inflammatory pain**. Acute inflammatory pain induced in male and female CD-1 mice as described in the Methods and in **Figure 2**. At the end of the terpene time course, hindpaw skin from the ipsilateral (LPS-treated) and contralateral (untreated) sides was collected and subjected to qPCR analysis for cytokine expression as noted. The data for each ipsilateral cytokine set was normalized to the value from the contralateral side (1.0 fold). Males and females for α-humulene treatment were separated for analysis based on an apparent difference in their response. Data shown as the mean ± SEM of 2 technical replicates, with sample sizes noted in each graph. *, ** = $p < 0.05, 0.01$ vs. same set vehicle group by Unpaired 2-Tailed *t* Test. Some terpenes appear to enhance expression of the anti-inflammatory cytokine IL-10, although geraniol also upregulated IL-6.

**Tables**

**Table S1:**

| Terpene | Chemical Structure (SMILES) | Mol. Weight | LogS | ASA_H | ASA_P |
|---|---|---|---|---|---|
| Geraniol | OC/C=C(/CC/C=C(\C)/C)\C | 154.3 | -2.4 | 314.1 | 87.4 |
| Linalool | OC(C=C)(CC/C=C(\C)/C)C | 154.3 | -2.1 | 292.8 | 103.5 |
| β-Pinene | C=C1C2C(C)C(C2)CC1 | 122.2 | -3.4 | 267.7 | 56.5 |
| α-Humulene | C/C/1=C\CC/C(/C)=C/CC(C)/C=C/C\1 | 190.3 | -3.4 | 357.2 | 57.1 |
| β-Caryophyllene | C=C1C2C(C(C)(C)C2)CCC(C)=CCC1 | 204.4 | -5.2 | 354.3 | 70.2 |

**Table S1: Physicochemical parameters of the modeled terpenes.** The chemical structure (SMILES), molecular weight, LogS, total hydrophobic surface area (ASA_H), and total polar surface area (ASA_P) are shown for each terpene. These were used as parameters in the molecular modeling experiments.
